# LZTR1 inactivation promotes MAPK/ ERK pathway activation in glioblastoma by stabilizing oncoprotein RIT1

**DOI:** 10.1101/2020.03.14.989954

**Authors:** Yuqi Wang, Jianong Zhang, Pingzhao Zhang, Zhipeng Zhao, Qilin Huang, Dapeng Yun, Juxiang Chen, Hongyan Chen, Chenji Wang, Daru Lu

**Author notes:** These authors contributed equally to this work (Yuqi Wang, Jianong Zhang, Pingzhao Zhang, Zhipeng Zhao). Corresponding Author: Daru Lu,; Chenji Wang,.

## Abstract

Large-scale sequencing studies on glioblastoma have identified numerous genetic alterations. Leucine-zipper-like transcription regulator 1 (*LZTR1*) is inactivated by non-synonymous mutations and copy number losses, suggesting that it is a tumor suppressor in glioblastoma. However, how *LZTR1* mutations contribute to glioblastoma pathogenesis remains poorly understood. Here, we revealed that LZTR1, as an adaptor of the CUL3 E3 ubiquitin ligase complex, recognizes and triggers ubiquitin-dependent degradation of oncoprotein RIT1, a RAS-like GTPase. Wild-type LZTR1 suppresses glioblastoma cell proliferation and migration by inactivating the MAPK/ERK signaling pathway in a RIT1-dependent manner. However, the effects were abrogated by the glioblastoma-associated LZTR1 mutations. Our findings revealed the underlying molecular mechanism of LZTR1 mutations-driven glioblastoma, and provide novel therapeutic target for LZTR1 mutations-driven glioblastoma.

## INTRODUCTION

Glioblastoma is the most common and deadliest form of brain tumor in adults; it is characterized by poor survival and high tumor heterogeneity (1–7). Despite the development of surgical techniques and chemoradiotherapy, the median survival rate of patients with glioblastoma remains 12–15 months (8–11). Therefore, novel therapeutic strategies for patients with glioblastoma should to be studied. Mapping the genomic landscape of glioblastoma has produced comprehensive molecular classification of these tumors, which may ultimately serve to improve the diagnosis and treatment of patients with glioblastoma (12–15), but the biological consequences of most alterations are still poorly understood.

Leucine-zipper-like transcription regulator (LZTR1) is a tumor suppressor in glioblastoma, receiving 4.4% nonsynonymous mutation and 22.4% focal deletion in the coding region and the locus, respectively (15). LZTR1, encoding a protein characterized by KELCH-BTB-BACK-BTB-BACK domain architecture, is an adaptor of CUL3-containing E3 ligase complex and localized in the Golgi apparatus (15–17). This protein binds to the N-terminal domain of CUL3 via the BTB-BACK domain and to substrates via the KELCH domain (15). Mutations in LZTR1 are also identified in Noonan syndrome (18–20) and schwannomatosis (21–24). Noonan syndrome is the most common RASopathy caused by mutations in genes involved in the RAS/MAPK signaling pathway, whereas schwannomatosis is a rare neurofibromatosis. However, the exact role of LZTR1 mutations and focal deletions in the pathogenesis of glioblastoma remain unknown.

RIT1 (RAS-like without CAAX1) encodes a RAS-family small GTPase with a high sequence homology to other RAS subfamily members, including K, H, R, N-RAS (25). This protein participates in the MAPK/ERK signaling pathway activation and mediates various physiological processes, such as cell proliferation, survival, and differentiation (26–30). RIT1 is overexpressed in various cancers and its increased expression is usually correlated with poor prognosis (31, 32). Furthermore, somatic mutations in RIT1 are identified in several human cancers (33–36). Interestingly, gain-of-function mutations in RIT1 are also observed in Noonan syndrome (37–41). However, the role of RIT1 in glioblastoma remains unclear.

In this study, we demonstrate that LZTR1 suppresses glioblastoma cell proliferation and migration by inactivating the MAPK/ERK pathway via promoting the ubiquitination and degradation of RIT1, an oncoprotein overexpressed in glioblastoma. Moreover, this effect is abrogated by glioblastoma-associated LZTR1 mutations. Our results established LZTR1 acts as a tumor suppressor in glioblastoma. Inhibition of MAPK/ERK pathway by small molecular inhibitors may represent a promising therapeutic strategy for targeting LZTR1-mutated glioblastoma.

## RESULTS

### RIT1 is identified as a novel LZTR1 interactor

To investigate the cellular functions of LZTR1 and identify the molecular mediators of LZTR1 in glioblastoma, we isolated the LZTR1 complex by using tandem affinity purification methods and determined the proteins present in the complex through mass spectrometry (**Fig. 1A, 1B**). Subunits of the CRL3-LZTR1 complex (CUL3 and RBX1) were detected in the complex. In addition to the known binding partner of LZTR1, RIT1 was co-purified in the LZTR1 complex (**Fig. 1B**). Considering that the gain-of-function mutations in RIT1 and the loss-of-function mutations in LZTR1 are present in Noonan syndrome, we investigated whether RIT1 was substrate of LZTR1.

**Figure 1.**
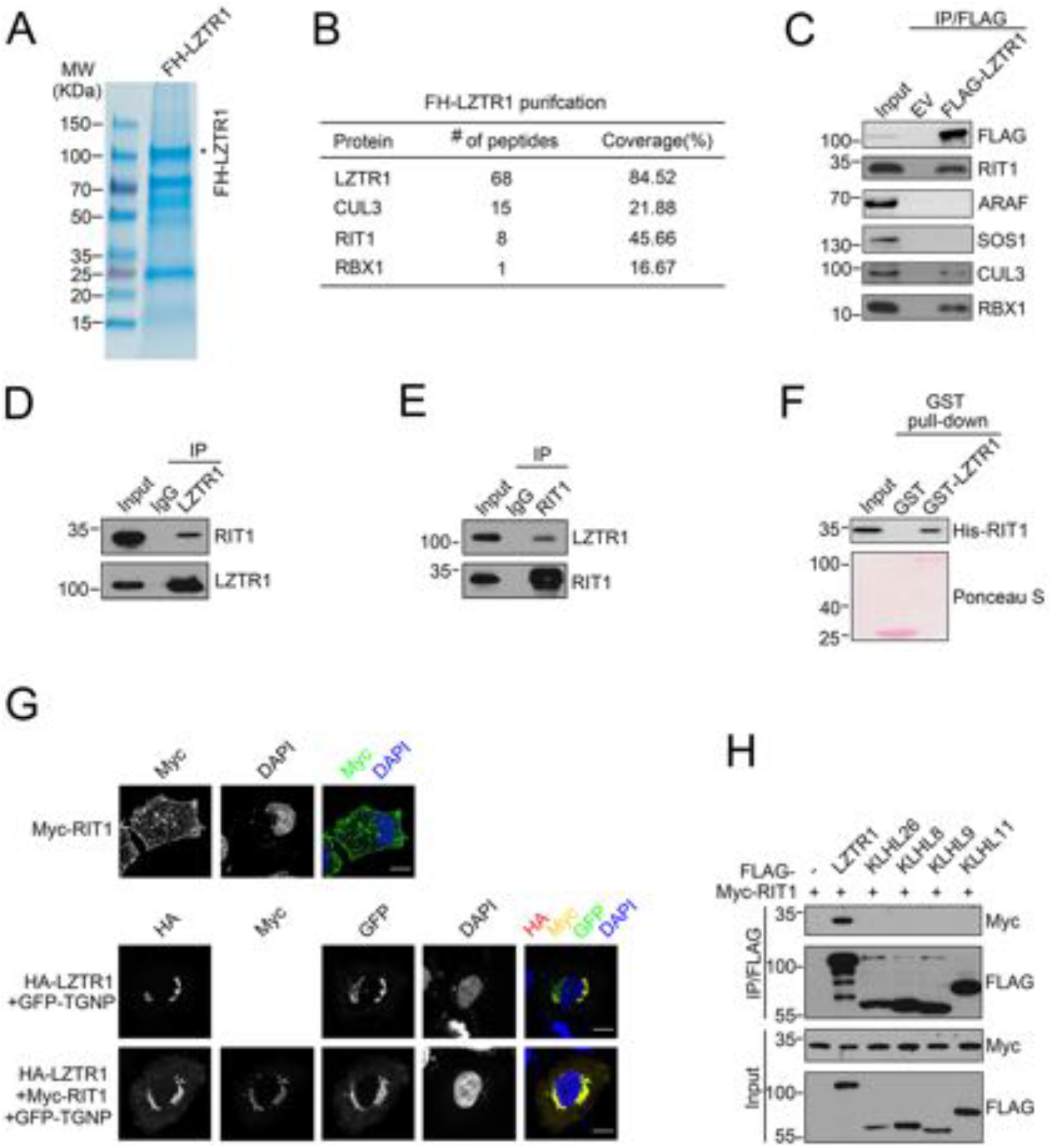
LZTR1 interacts with RIT1. (A, B) Tandem affinity purification of the LZTR1-containing protein complex was conducted using 293T cells stably overexpressing FLAG-HA(FH)-LZTR1. Associated proteins were separated by SDS-PAGE and visualized by Coomassie Blue staining (A). The number of peptides and coverage identified by mass spectrometry analysis is shown in the table (B). (C) Western blot of indicated protein in WCLs and co-IP samples of anti-FLAG antibody obtained from 293T cells transfected with indicated plasmids. (D) Western blot of indicated proteins in WCLs and co-IP samples of IgG or anti-LZTR1 antibody obtained from the cell extracts of U87-MG cells treated with 20 μM of MG132 for 8 h. (E) Western blot of indicated proteins in WCLs and co-IP samples of IgG or anti-RIT1 antibody obtained from the cell extracts of U87-MG cells treated with 20 μM of MG132 for 8 h. (F) Bacterially expressed GST-LZTR1 proteins or GST bound to glutathione-sepharose beads and incubated with bacterially expressed His-RIT1 proteins. Bound His-RIT1 proteins were detected by Western blotting with anti-His antibody, and GST fusion proteins were detected by Ponceau staining. (G) Representative immunofluorescence images of HeLa cells transfected with indicated plasmids, stained with RIT1(Myc), LZTR1(HA), TGNP(GFP) and DAPI. Scale bar, 20 μm. (H) Western blot of indicated proteins in WCLs and co-IP samples of anti-FLAG antibody obtained from 293T cells transfected with indicated plasmids.

To verify that RIT1 is a LZTR1-interacting protein, we examined whether LZTR1 interacted with RIT1 in cells. Co-immunoprecipitation (co-IP) analysis showed that, endogenous RIT1, but not the other members of the RAS/MAPK pathway (ARAF, SOS1), interact with FLAG-LZTR1 (**Fig. 1C**). Next, we extended our analysis by investigating whether endogenous LZTR1 and RIT1 can interact with each other. We performed co-IP by using LZTR1 or RIT1 antibody. As shown in **Fig. 1D, 1E**, endogenous LZTR1 protein was co-immunoprecipitated with RIT1 and endogenous RIT1 protein was co-immunoprecipitated with LZTR1. To determine whether the interaction between LZTR1 and RIT1 is direct, we detected their interaction in vitro by using purified recombinant proteins. As shown in **Fig. 1F**, GST-LZTR1, but not the GST alone, bound to His-RIT1 in a pull-down assay, suggesting LZTR1 and RIT1 physically interacted with each other.

To further examine the interaction between LZTR1 and RIT1 in vivo, we determined whether these two proteins are co-localized to the same subcellular compartments. Previous studies suggested LZTR1 is a Golgi apparatus-localized protein (16), we investigated the subcellular localization that LZTR1-RIT1 interaction occurs. As shown in **Fig. 1G**, the exogenous transfected RIT1 protein was detected at cytoplasm and plasma membrane, and the exogenous transfected LZTR1 protein was detected at Golgi apparatus. However, LZTR1 can recruit RIT1 to Golgi apparatus, as demonstrated the co-localization of RIT1 with Golgi apparatus marker TGNP. Similar results were obtained by using GM130 or Golgi-tracker Red dye to mark the Golgi apparatus (**Fig. S1**). Additionally, we found that only LZTR1, but not the other CUL3-based BTB-domain-containing adaptors (KLHL26, KLHL8, KLHL9 and KLHL11), interacted with RIT1 (**Fig. 1H**). Taken together, these data suggest that RIT1 is a LZTR1-interacting protein.

### LZTR1 promotes RIT1 degradation and ubiquitination

RIT1 plays a critical role in the RAS/MAPK pathway (26–30), but its functional regulation by post-translational modifications is poorly understood. The treatment of glioblastoma cell lines U87-MG or U118-MG with the proteasome inhibitors bortezomib or MG-132 inevitably increased RIT1 protein levels but not corresponding mRNA levels (**Fig. 2A**, **Fig. S2A**). MLN4924, a small-molecule inhibitor of NEDD8-activating enzyme required for CRL complex activation, also caused the accumulation of RIT1 proteins but not of mRNA (**Fig. 2A**, **Fig. S2A**). We depleted RBX1 or each Cullin adaptor, including CUL1, CUL2, CUL3, CUL4A, CUL4B, and CUL5, in U87-MG cells by small interfering RNAs. Only RBX1 or CUL3 depletion led to a marked increase in the abundance of RIT1 proteins, but not of mRNA, suggesting that RIT1 protein stability may be regulated by a CUL3-RBX1 ubiquitin ligase complex(s) (**Fig. 2B**, **Fig. S2B**). Furthermore, RIT1 protein was elevated in dominant-negative fragment of CUL3 (CUL3-DN) overexpressed cells, but this phenomenon was not observed in CUL1, 2, 4A, 4B, 5-DN - overexpressed cells (**Fig. 2C**). We found that the overexpression of LZTR1, but not the other examined CUL3-based adaptors, markedly decreased RIT1 protein levels (**Fig. 2D**). The degradative effect is completely reversed by treatment with MG132, bortezomib, or MLN4924 (**Fig. 2E**). Moreover, only RIT1, but not the other examined components of the RAS/MAPK pathway, is degraded by LZTR1 (**Fig. 2F**). Moreover, only RIT1, but not the other examined members of RAS/MAPK pathway, is downregulated in LZTR1-overexpressed 293T cells (**Fig 2G**). Conversely, only RIT1, but not the other examined members of RAS/MAPK pathway, is upregulated in LZTR1-depleted U87-MG or U118-MG cells (**Fig. 2H**). LZTR1 depletion had no overt effect on corresponding RIT1 mRNA levels in U87-MG or U118-MG cells (**Fig. S2C**). In the absence of de novo protein synthesis, the half-life of endogenous RIT1 protein is considerably longer in the LZTR1-depleted cells than that in the control cells (**Fig. 2I, 2J**). As shown in **Fig. 2K**, overexpression of LZTR1, but not the other examined CUL3-based adaptors, enhanced RIT1 polyubiquitination. Overall, these data demonstrated that the LZTR1-CUL3-RBX1 E3 ubiquitination ligase complex regulates RIT1 protein stability through ubiquitin-dependent proteasomal degradation in glioblastoma cells.

**Figure 2.**
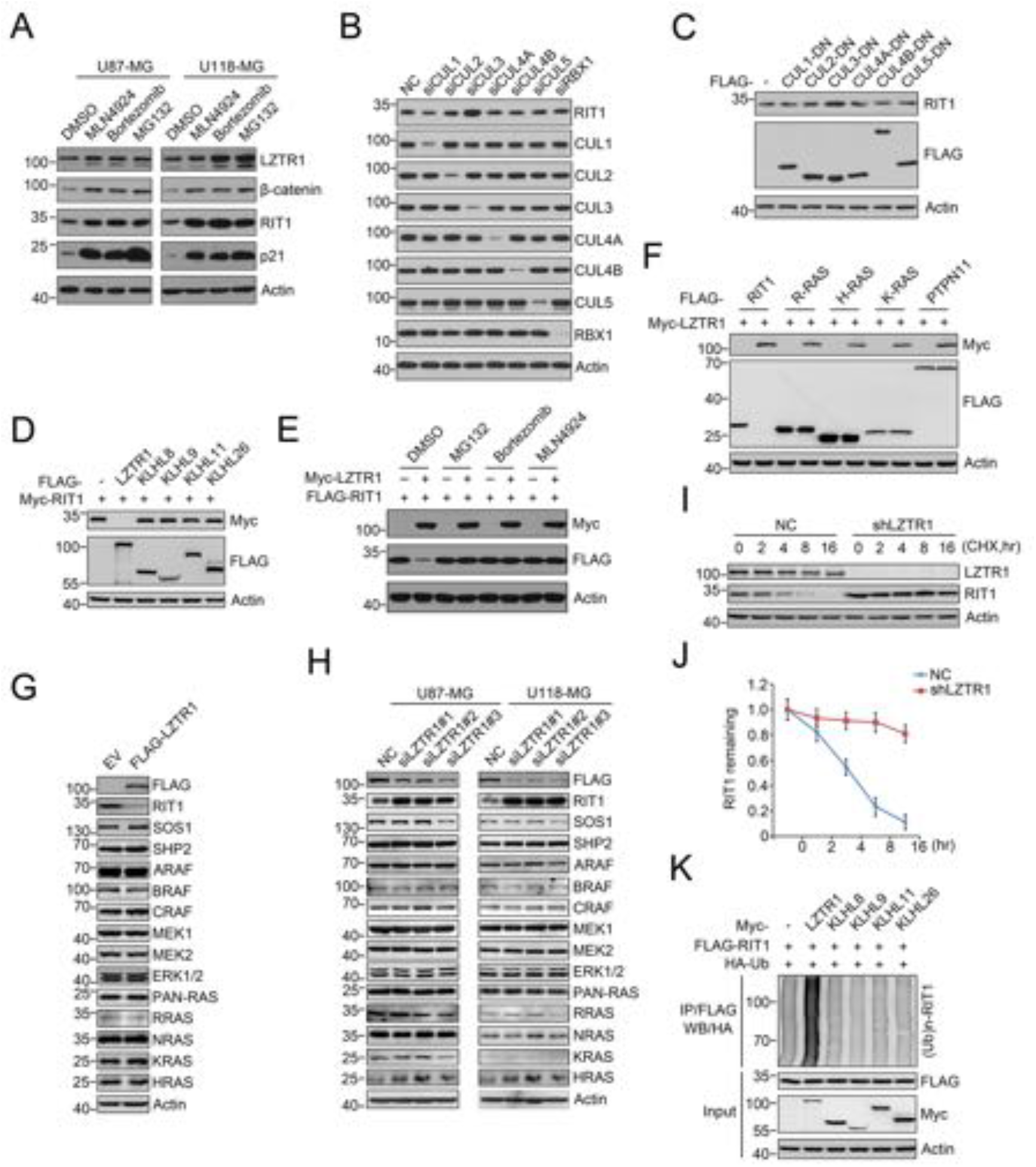
LZTR1 promotes RIT1 degradation and ubiquitination. (A) Western blot of indicated proteins in WCLs from U87-MG or U118-MG cells treated with DMSO, MLN4924(100 nM), Bortezomib (20 nM) or MG132 (20 μM) for 8h. (B) Western blot of indicated proteins in WCLs from U87-MG cells transfected with indicated siRNAs. (C, D) Western blot of indicated proteins in WCLs from 293T cells transfected with indicated plasmids. (E) Western blot of indicated proteins in WCLs from 293T cells transfected with indicated plasmids with DMSO, MG132 (20 μM), Bortezomib (20 nM) or MLN4924(100 nM) for 8 h. (F) Western blot of indicated proteins in WCLs from 293T cells transfected with indicated plasmids. (G) Western blot of indicated proteins in WCLs from U87-MG cells infected with lentivirus expressing EV or FLAG-LZTR1 for 72 h. (H) Western blot of indicated proteins in WCLs from U87-MG or U118-MG cells transfected with negative control (NC) or LZTR1-specific siRNAs. (I, J) Western blot of indicated proteins in WCLs of U87-MG cells infected with lentivirus expressing LZTR1-specific shRNA or NC for 72 h and then treated with 50 μg/ml cycloheximide and harvested at different time points (I). At each time point, the intensity of RIT1 was normalized to the intensity of actin and then to the value at 0 h (J). Similar results were obtained from three independent experiments. (K) Western blot of the products of in vivo ubiquitination assays from 293T cells transfected with the indicated plasmids and treated with 20 μM MG132 for 8 h.

### Identification the mutual interaction domain between RIT1 and LZTR1

To determine which domain of LZTR1 was required for RIT1 binding, we generated a panel of LZTR1 truncation mutants (**Fig. 3A**). A co-IP assay demonstrated that the RIT1 binding capacity of LZTR1 was totally abolished by deletion of the KELCH domain, but not the BTB or BACK domains, suggesting that the KELCH domain of LZTR1 is required for RIT1 binding (**Fig. 3B**). All the truncation mutants of LZTR1 lost the capacity to promote RIT1 degradation (**Fig. 3C**) and polyubiquitination (**Fig. 3D**), suggesting the KELCH, BTB and BACK domains are indispensable for LZTR1-mediated RIT1 destruction. Moreover, we found that the two BTB-BACK domains are required for LZTR1 dimerization (**Fig. 3E**). To determine which domain of RIT1 was required for LZTR1 binding, we generated a panel of RIT1 truncation mutants (**Fig. 3F**). The co-IP assay demonstrated that the region corresponding the 1-180 aa of RIT1 is required for LZTR1 binding (**Fig. 3G**). Therefore, our findings demonstrated that the KELCH domain of LZTR1 and the region corresponding the 1-180aa of RIT1 mediated their interaction.

**Figure 3.**
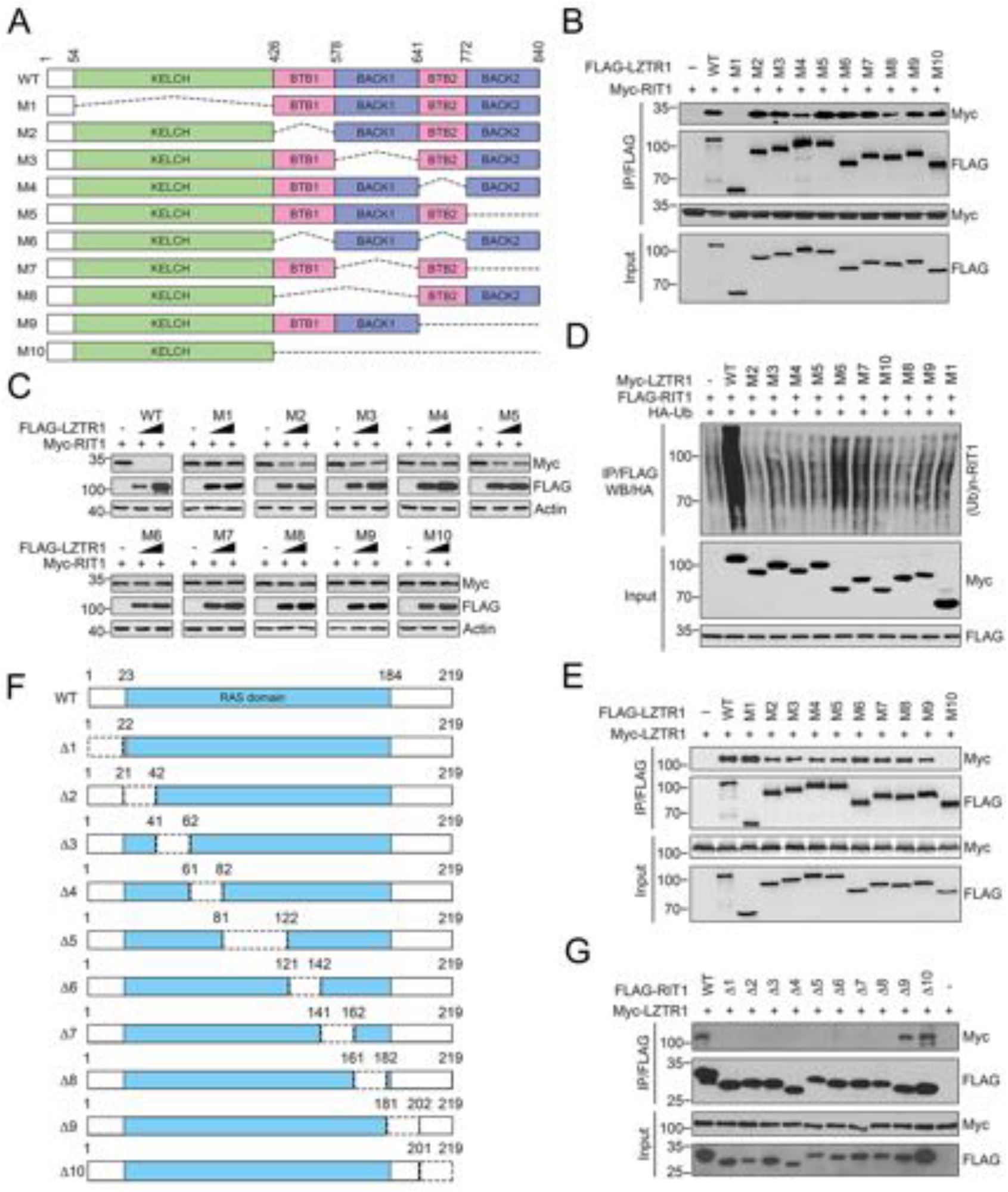
Identification of the mutual-binding regions of LZTR1 and RIT1. (A) Diagram showing the full-length LZTR1 and its deletion mutants (M1-M10). (B) Western blot of indicated proteins in WCLs and samples from co-IP with anti-FLAG antibody in 293T cells transfected with the indicated plasmids. (C) Western blot of indicated proteins in WCLs from 293T cells transfected with the indicated plasmids. (D) Western blot of the products of in vivo ubiquitination assays from 293T cells transfected with the indicated plasmids and treated with 20 μM MG132 for 8 h. (E) Western blot of indicated proteins in WCLs and samples from co-IP with anti-FLAG antibody in 293T cells transfected with the indicated plasmids. (F) Diagram showing the full-length RIT1 and ten RIT1 splicing variants (Δ1-Δ10). (G) Western blot of indicated proteins in WCLs and samples from co-IP with anti-FLAG antibody in 293T cells transfected with the indicated plasmids.

### Glioblastoma-associated mutants of LZTR1 are defective in promoting RIT1 degradation and ubiquitination

Non-synonymous mutations with loss of heterozygosity in LZTR1 was present a subset of glioblastoma patients (15). We hypothesized that glioblastoma–associated LZTR1 mutants would impair their ability to degrade RIT1 proteins. We generated a series of glioblastoma-associated LZTR1 mutants, and examined their interactions with RIT1 via co-IP assay. Among the ten LZTR1 mutations, nine occur in the KELCH domain, which is responsible for RIT1 binding (**Fig. 4A**). As shown in **Fig 4B**, except R810W, the RIT1-binding ability of LZTR1 mutants was greatly impaired compared with that of wild-type LZTR1. The LZTR1-mediated degradation and ubiquitination of RIT1 were abolished or markedly attenuated for these mutants (**Fig. 4C, 4D**). R810W mutation was in the second BACK domain of LZTR1 which was not required for RIT1 binding, but this mutation may disrupt LZTR1 dimerization or CUL3 binding. Moreover, we found that Noonan syndrome and schwannoma-associated LZTR1 mutants were also defective in promoting RIT1 degradation and ubiquitination (**Supplementary Fig. 3A-E**). Overall, our findings suggested that RIT1 stability may be dysregulated by glioblastoma and other LZTR1 mutation-related human diseases.

**Figure 4.**
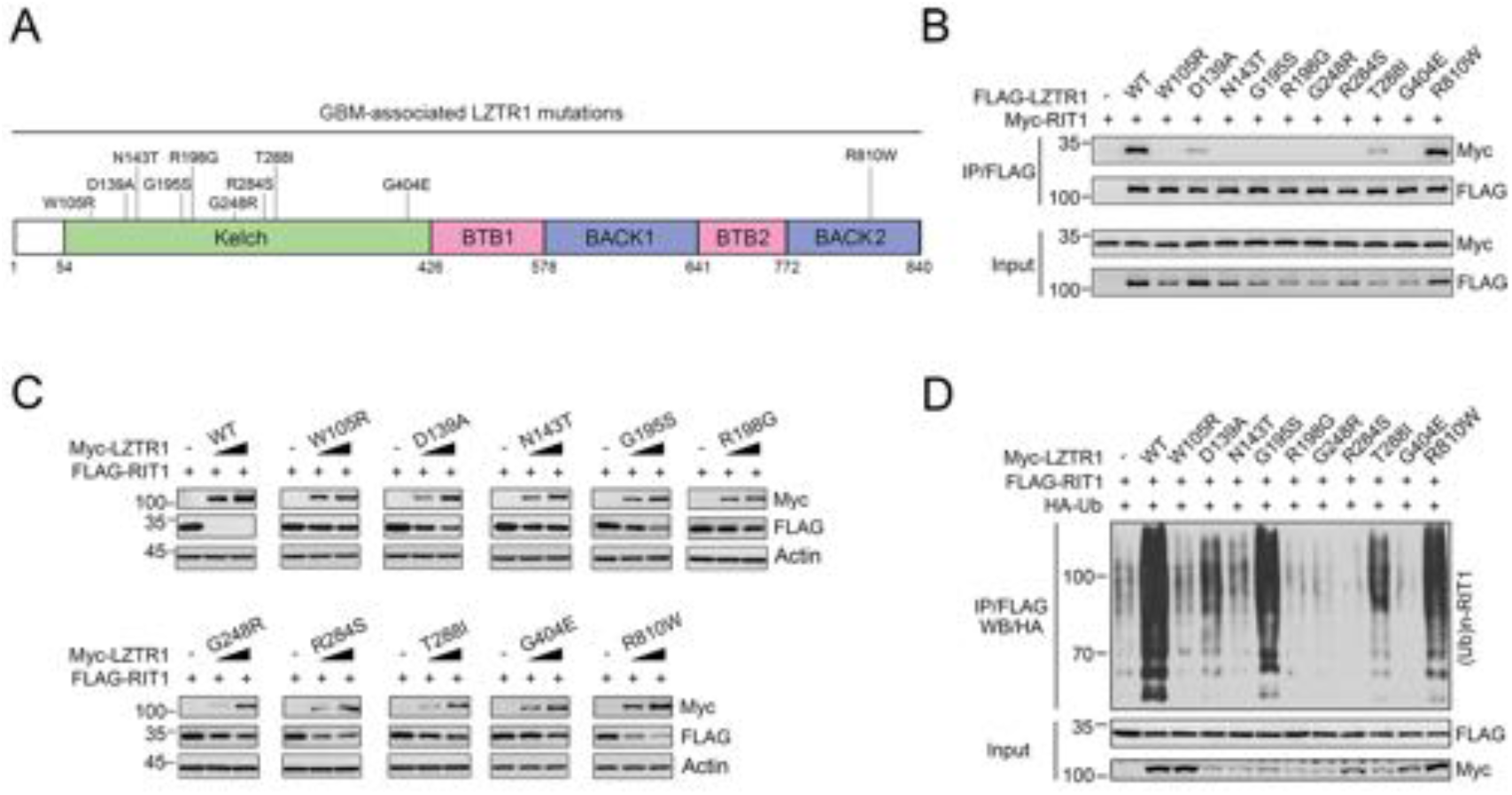
Glioblastoma-associated LZTR1 mutants are defective in promoting RIT1 degradation and ubiquitination. (A) Diagram showing the glioblastoma-associated LZTR1 mutations. (B) Western blot of indicated proteins in WCLs and samples from co-IP with anti-FLAG antibody in 293T cells transfected with the indicated plasmids and treated with 20 μM MG132 for 8 h. (C) Western blot of indicated proteins in WCLs from 293T cells transfected with the indicated plasmids. (D) Western blot of the products of in vivo ubiquitination assays from 293T cells transfected with the indicated plasmids and treated with 20 μM MG132 for 8 h.

### LZTR1 suppresses MAPK/ERK pathway activation partly through RIT1

RIT1 plays a critical role in regulating MAPK/ERK pathway activation (26–30). We examined the MAPK/ERK activation in LZTR1-or RIT1-knocked out (KO) glioblastoma cells established by CRISPR-Cas9 methods. We observed a decreased MAPK/ERK pathway activation in EGF-stimulated RIT1-KO cells than that in parental cells (**Fig. 5A, 5B**). Conversely, we observed a stronger MAPK/ERK activation in EGF-stimulated LZTR1-KO cells, and the phenotype could be rescued upon co-knockout of RIT1 or overexpression of wild-type LZTR1, but not the glioblastoma-associated LZTR1-G248R mutant (**Fig. 5A, 5B**). Moreover, we observed an increased MAPK/ERK response to EGF stimulation in RIT1-overexpressing cells (**Fig. 5C, 5D**). Overall, our findings suggest that LZTR1 modulates MAPK/ERK pathway activation, at least in part, depending on its substrate RIT1.

**Figure 5.**
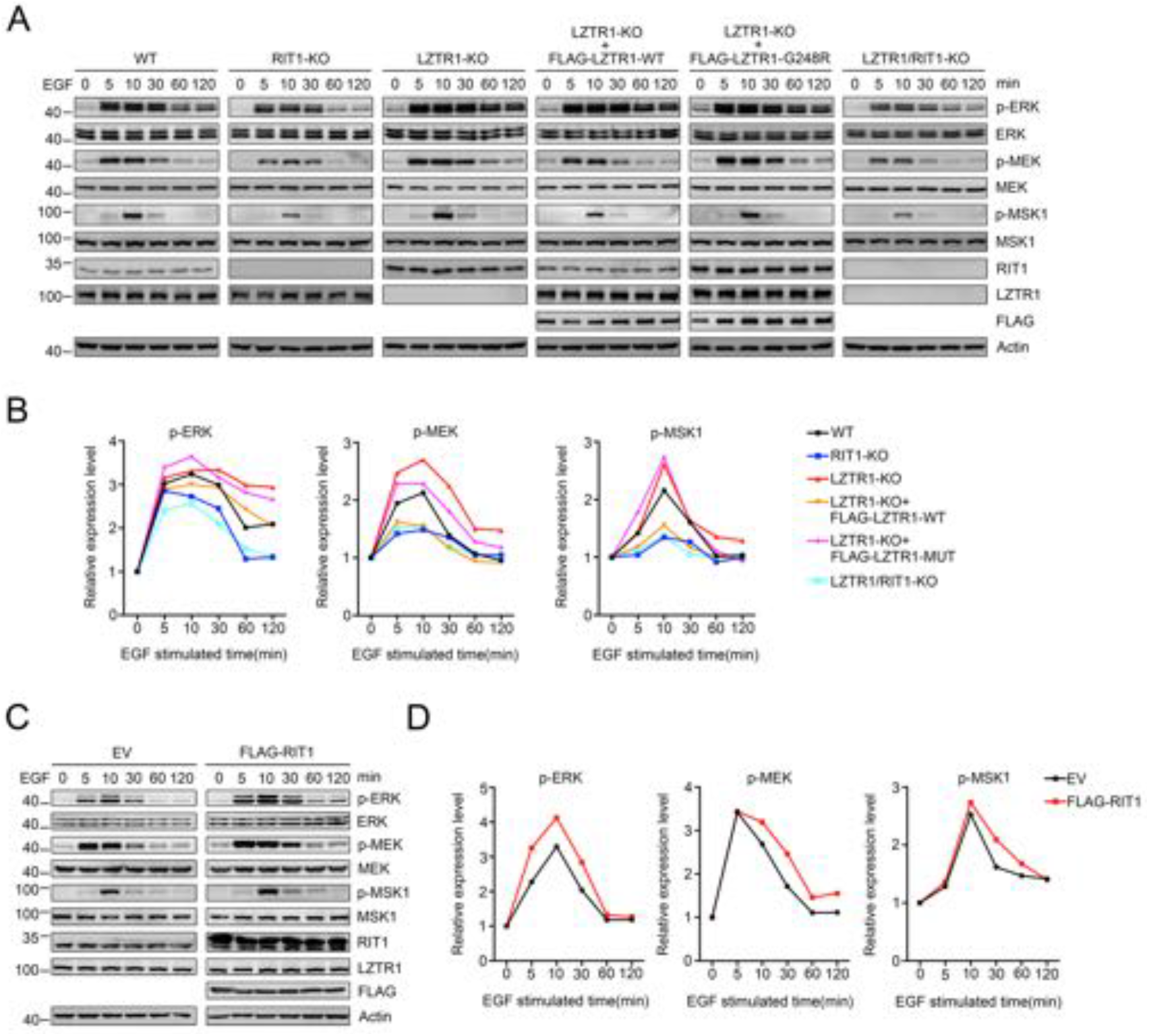
LZTR1 suppresses MAPK/ERK pathway through RIT1 destruction. (A) Western blot of indicated proteins in WCLs from U87-MG cells (WT or LZTR1/RIT1 KO) infected with lentivirus expressing, treated with 50 ng/ml EGF after precultured in serum-free medium for 48h and harvested at different time points. (B) At each time point, the intensity of p-ERK, p-MEK, p-MSK1 were normalized to the intensity of ERK, MEK, MSK1 respectively and then to the value at 0 min. (C) Western blot of indicated proteins in WCLs from U87-MG cells infected with lentivirus expressing indicated plasmids, treated with 50 ng/ml EGF after precultured in serum-free medium for 48h and harvested at different time points. (D) At each time point, the intensity of p-ERK, p-MEK, p-MSK1 were normalized to the intensity of ERK, MEK, MSK1 respectively and then to the value at 0 min.

### LZTR1 suppresses glioblastoma cell growth in a RIT1-dependent manner

To further address the biological importance of LZTR1-RIT1 axis in glioblastoma, we stably depleted their expression respectively or simultaneously in U87-MG cells (**Fig. 6A**). As determined by the CCK-8 assay, the growth rate of RIT1-depleted U87-MG cells was slower than that of the control cells, whereas depletion of LZTR1 could markedly promote cell growth, and the phenotypes could be rescued by RIT1 co-depletion (**Fig. 6B**). Stably overexpression of RIT1 markedly promoted cell proliferation. In contrast, stably overexpression of LZTR1, but not the glioblastoma-associated LZTR1 mutants (W105R,G248R), markedly decreased cell proliferation (**Fig. S4A**). Similarly, we found LZTR1 depletion had a positive effect, whereas RIT1 depletion had a negative effect on cell cycle (**Fig. 6C**), DNA synthesis (**Fig. 6D, 6F**) and cell migration (**Fig. 6E, 6G**). Moreover, LZTR1 depletion-caused cell phenotype changes can be reversed by RIT1 co-depletion (**Fig. 6B-G**). The opposing roles of LZTR1 and RIT1 in cell cycle, DNA synthesis and cell migration were also demonstrated in LZTR1- or RIT1- overexpressing cells (**Fig. S4B-F**).

**Figure 6.**
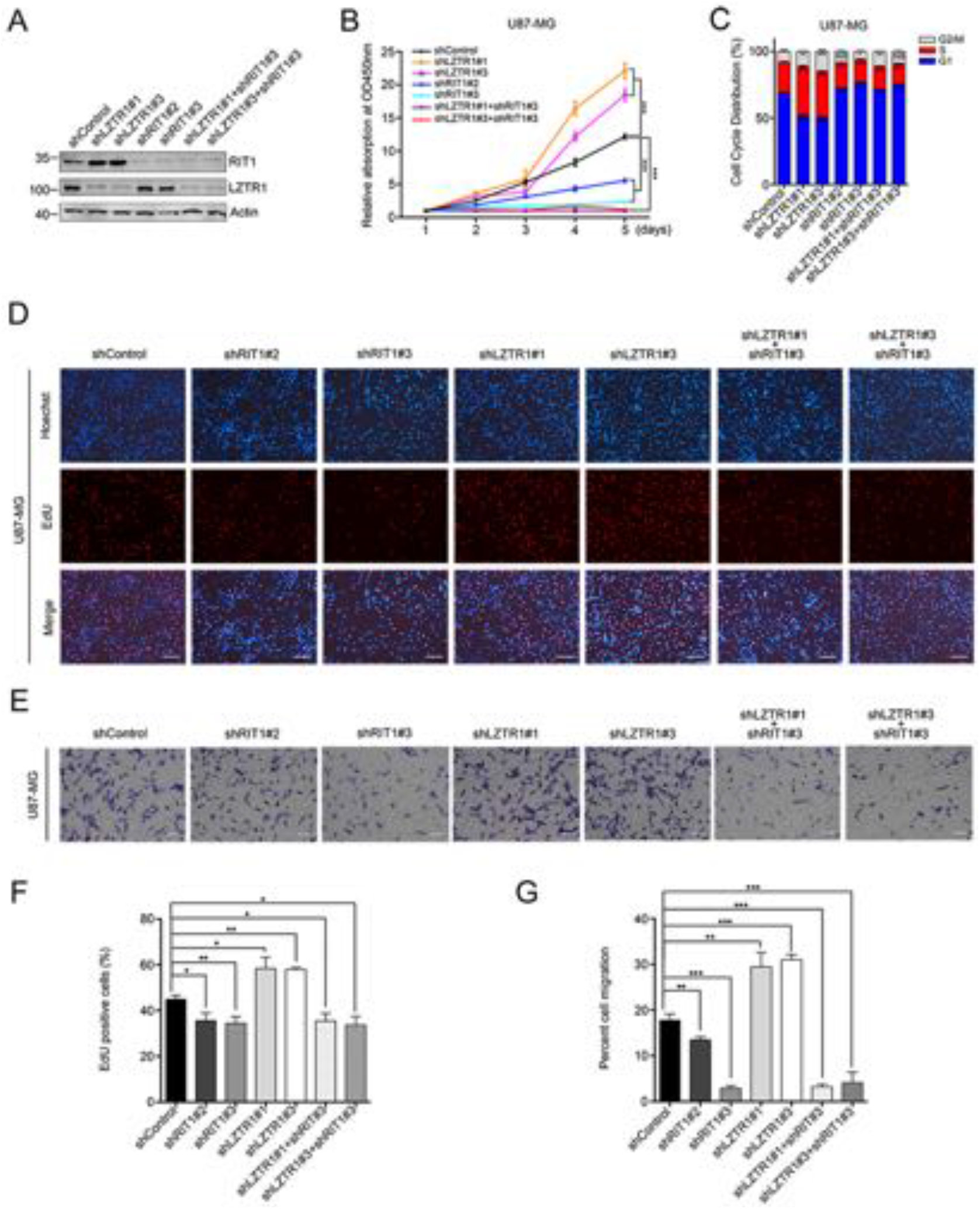
LZTR1 suppresses U87-MG cell growth and migration in a RIT1-dependent manner. (A) Western blot of indicated proteins in WCLs of U87-MG cells infected with lentivirus expressing indicated shRNAs for 72 h. (B) CCK-8 cell proliferation analysis of U87-MG cells infected with lentivirus expressing indicated shRNAs. Data are shown as means ± SD (n=3). ***p<0.001. (C) Cell cycle analysis of U87-MG cells infected with lentivirus expressing indicated shRNAs. Data are shown as means ± SD (n=3). *p<0.05, **p<0.01, ***p<0.001. (D) EdU incorporation analysis of U87-MG cells infected with lentivirus expression indicated shRNAs. Scale bar, 20 μm. (E) Cell migration analysis of U87-MG cells infected with lentivirus expression indicated shRNAs. Scale bar, 20 μm. (F, G) The quantitative analysis of EdU incorporation assay (F), and cell migration assay (G). Data are shown as means ± SD (n=3). *p<0.05, **p<0.01, ***p<0.001.

Our data demonstrated that LZTR1 regulated RIT1-dependent glioblastoma cell growth *in vitro*. We then performed *in vivo* experiments with intracranial xenotransplanted tumor models. In intracranial xenograft mice, tumor progression was monitored by bioluminescence imaging on days 14, 20 and 26 after implantation. The mice bearing RIT1-overexpressing or LZTR1-depleted U87-MG xenografts showed markedly increases in tumor growth compared with those in the control groups (**Fig. 7A**). In contrast, the mice bearing RIT1-depleted or LZTR1-overexpressing xenografts showed noticeable regression of tumor growth compared with those in the control groups (**Fig. 7A**). The mice bearing LZTR1/RIT1 co-depleted U87-MG xenografts displayed phenotypes like that of the RIT1-depleted group (**Fig. 7A**). The survival rates of different groups of intracranial xenograft mice were described by the survival curve. As shown in **Fig. 7B**, the OS (overall survival) time of the RIT1-overexpressing and LZTR1-depleted groups were shorter than that of the control mice. The OS time of the LZTR1-overexpressing, RIT1-depleted and LZTR1/RIT1 co-depleted groups had longer OS time than that of the control groups. The representative images of the paralleled HE-stained tumor cytostructure are shown in **Fig. 7C**. Immunostaining analysis revealed an increase in Ki-67 and phospho-ERK expression levels in tumors derived from RIT1-overexpressing or LZTR1-depleted U87-MG xenografts (**Fig. 7D, 7E**). Taken together, our data suggests that LZTR1 negatively regulates glioblastoma cell growth *in vitro* and *in vivo* at least in part, dependent on RIT1.

**Figure 7.**
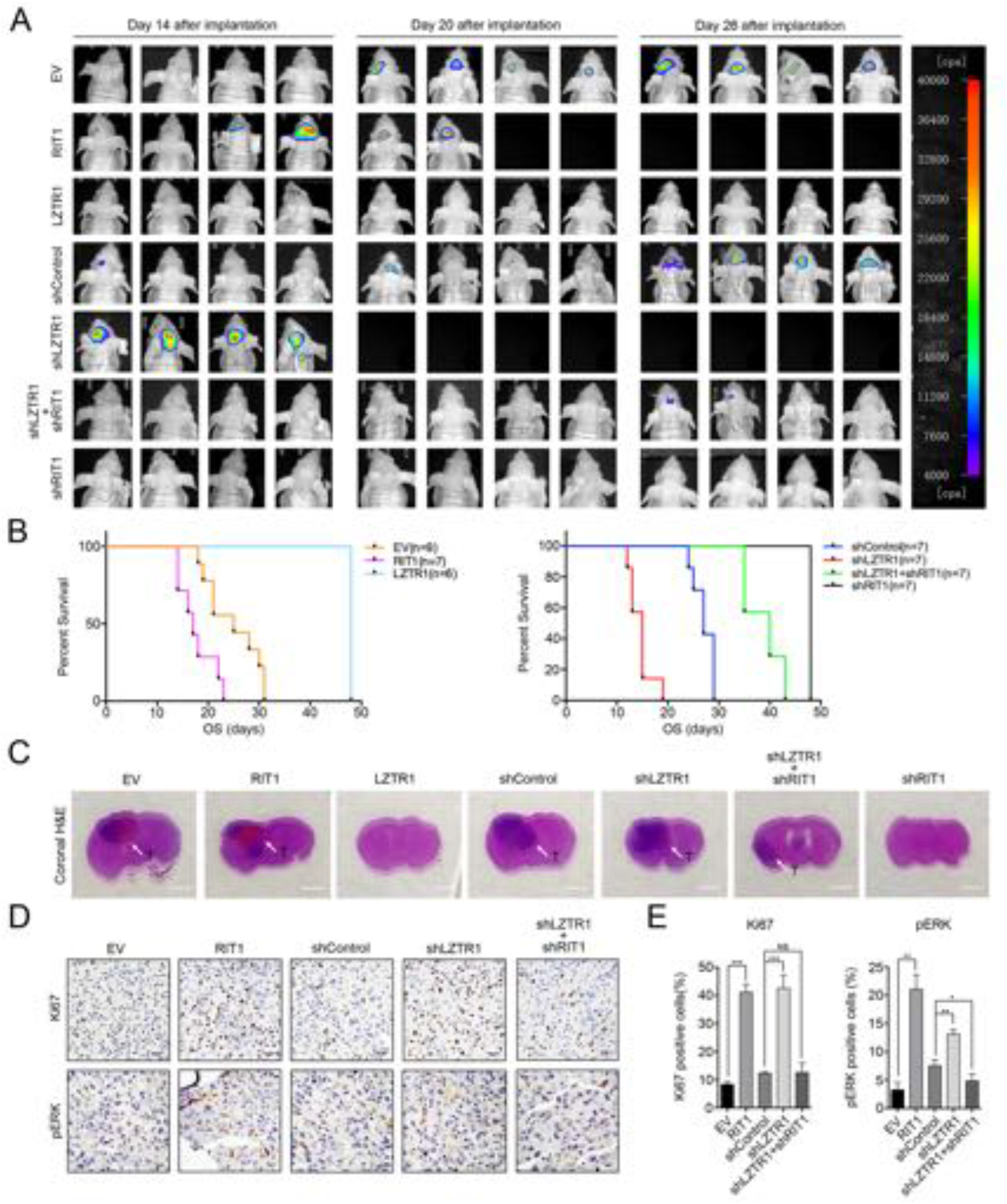
LZTR1 suppresses intracranial glioblastoma xenografts growth in a RIT1-dependent manner. (A) Representative pseudocolor bioluminescence images of intracranial mice of U87-MG cells with lentivirus expression indicated proteins or shRNAs on the days as indicated. (B) Kaplan-Meier curve determining the survival rate of mice with intracranial xenografts derived from U87-MG cells with lentivirus expression indicated proteins or shRNAs. (C) Representative H&E staining for a pathological form of the intracranial tumor from each group; T, tumor. (D, E) IHC analysis of Ki67 and p-ERK expression in intracranial tumors from U87-MG cells with lentivirus expression indicated proteins shRNAs. Data are shown as means ± SD (n=3). *p<0.05, **p<0.01, ***p<0.001.

### RIT1 is increased in glioma and serves as a prognostic factor

To investigate the expression of RIT1 in glioma, we analyzed the mRNA expression of RIT1 in three independent cohorts, including TCGA, CGGA, and Rembrandt. As shown in **Fig. 8A**, the mRNA expression of RIT1 increased in human gliomas in accordance with WHO grades. To investigate the relationship between RIT1 expression and clinical prognosis, we examined the prognostic significance of RIT1 expression in the TCGA and CGGA cohorts. Kaplan-Meier analyses showed that RIT1 expression did not affect the clinical outcome of the patients with gliomas in the TCGA cohort (**Fig. 8B**). However, the high RIT1 mRNA expression could significantly predict unfavorable OS and PFS for the patients with gliomas in the CGGA cohort (**Fig. 8B**). Considering that a higher tumor grade is an indubitable risk factor for gliomas, we further obtained similar results in patients with high-grade gliomas (**Fig. 8B**). Taken together, these results indicated that RIT1 could be a prognostic factor of human glioma.

**Figure 8.**
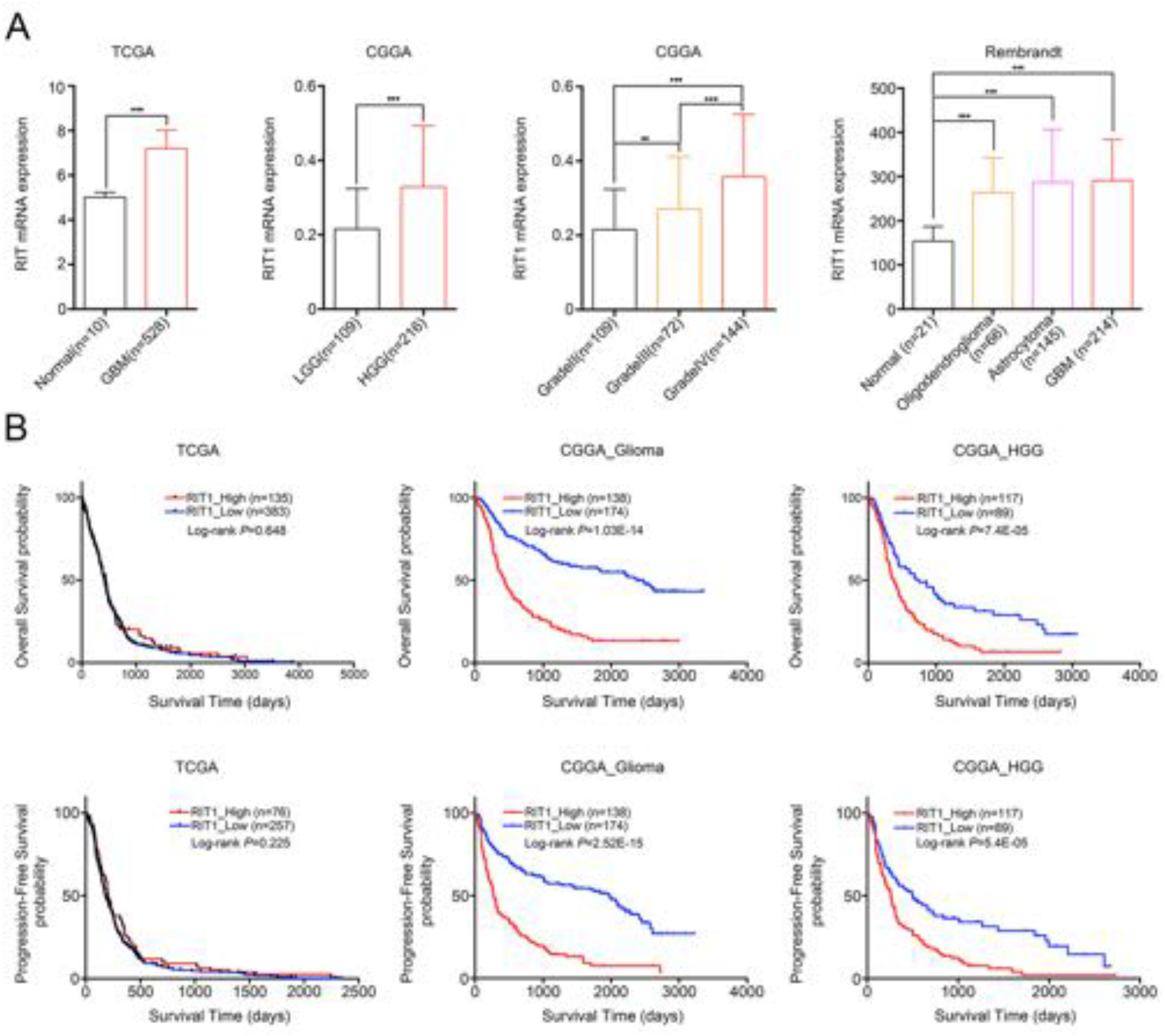
High expression of RIT1 predicts a poor clinical outcome in human glioblastomas. (A) Analysis of RIT1 mRNA expression level in TCGA, CGGA and Rembrandt cohorts. Data are shown as means ± SD. **p<0.01, ***p<0.001. (B) Kaplan-Meier curves were plotted according to different RIT1 mRNA expression for overall survival (upper panels) and progression-free survival (lower panels) in TCGA (left panels) and CGGA (right panels) cohorts.

## Discussion

Although mutations with the loss of *LZTR1* heterozygosity were observed in glioblastoma (15), the mechanisms of pathogenesis remain poorly understood. Previous studies demonstrated that LZTR1 encodes an adaptor of CUL3-containing E3 ligase complexes, and some glioblastoma-associated LZTR1 mutations occurred in the BTB or BACK domain that perturbed their interactions with CUL3 (15). Moreover, the loss-of-function mutations of LZTR1 promoted the self-renewal of cancer stem cells and growth of glioblastoma spheres (15). In the current study, RIT1 was identified as a novel substrate of LZTR1-CUL3-RBX1 ubiquitin ligase complex that plays crucial roles in glioblastoma cell growth *in vitro* and *in vivo*. LZTR1 inactivates the MAPK/ERK pathway and suppresses cell proliferation and migration in glioblastoma by targeting RIT1 for ubiquitin-dependent degradation, but this effect is abrogated by glioblastoma-associated LZTR1 mutations. Collectively, we identified a novel LZTR1-mutation-driven protumorigenic process in glioblastoma by activating the MAPK/ERK pathway (**Fig. 9**). Elucidation of this underlying molecular mechanism would be helpful to identify patients benefitting from MAPK/ERK pathway inhibitors and develop novel therapeutic approaches for patients carrying LZTR1 mutations.

**Figure 9.**
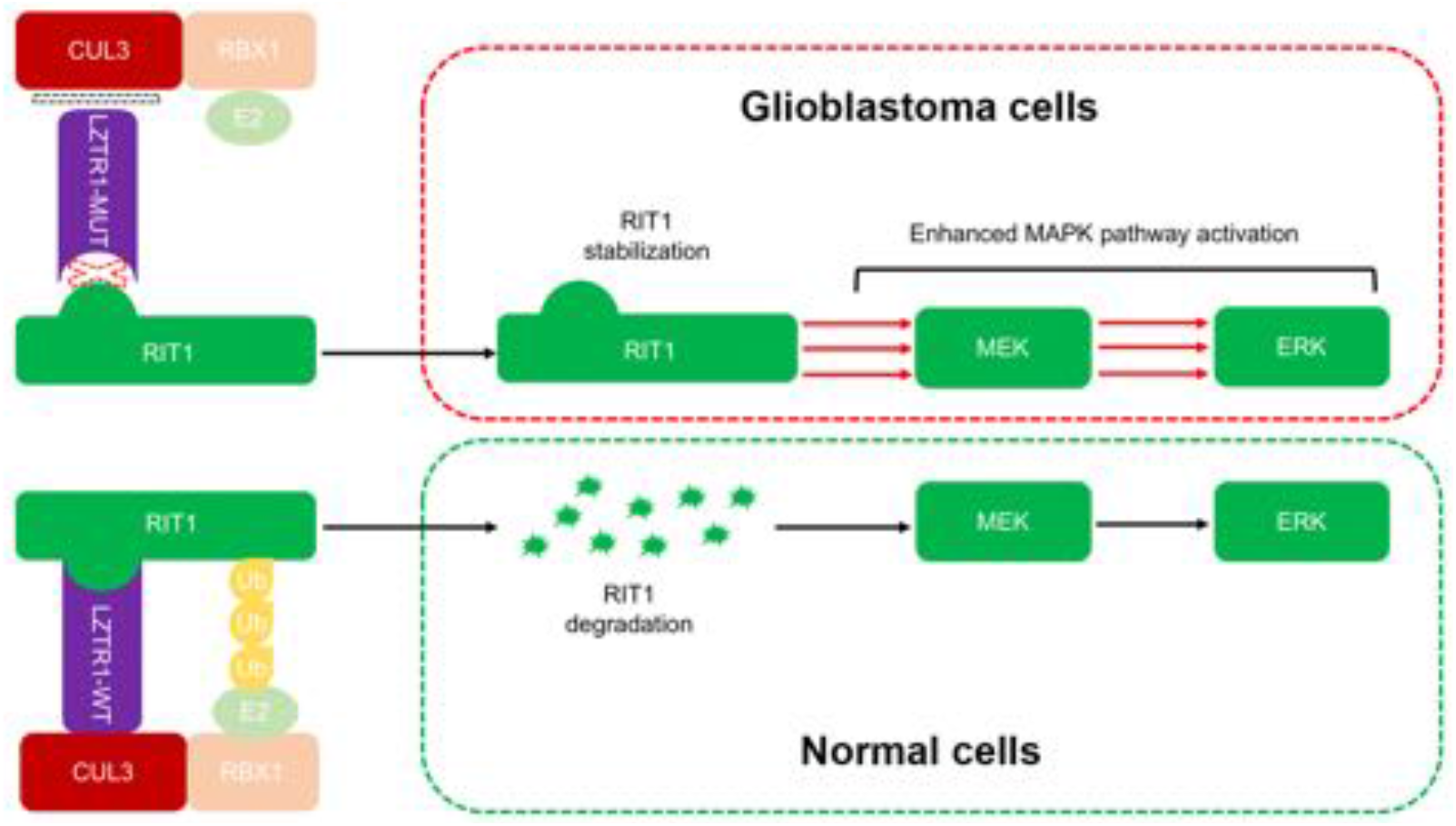
Schematic of the proposed mechanism for LZTR1-mediated RIT1 ubiquitination and enhanced MAPK/ERK pathway activation by LZTR1 inactivation.

Genetic studies have mapped more than 50 different mutations to *LZTR1* in various diseases (15, 18, 22, 42–44). As an adaptor of CUL3-containing E3 ligase complex, identification of LZTR1 substrates would provide functional insights into the pathogenesis of glioblastoma and other LZTR1-related diseases. However, very limited number of LZTR1 substrates had been identified and functionally explored. A recent study reported that LZTR1 promoted the nonproteolytic ubiquitination of K, N, R-RAS and hyperactivate the RAS/MAPK signaling pathway (45, 46). During our manuscript preparation, Castel et al. also reported that LZTR1 regulates the ubiquitination and degradation of RIT1 (47). Although most of biochemical results from that paper, such as protein interaction, protein degradation, are consistent with our study, the biological importance of LZTR1-RIT1 axis dysregulation in glioblastoma was still unclear (47). Thus, these studies together with ours, suggested LZTR1 could suppress MAPK/ERK pathway activation through promoting the proteolytic or nonproteolytic ubiquitination of multiple components of this pathway.

RIT1 is reported to be overexpressed or constitutively mutated in multiple cancers (31–36). However, the role of RIT1 in glioblastoma remains unknown. In our study, we demonstrated that the RIT protein stability was elevated in LZTR1-inacviated glioblastoma cells. Additionally, we also showed the mRNA levels of RIT1 was elevated, and its high expression predicts a poor clinical outcome in glioblastoma (**Fig. 8**). Previous studies demonstrated that RIT1 is involved in various biological processes, including cell proliferation, migration, cell survival, and neuronal morphogenesis (26–30). In our study, we demonstrated that RIT1 plays a positive role in glioblastoma cell growth *in vitro* and *in vivo* by enhancing MAPK/ERK activation. Therefore, the aberrant activation of RIT1 could be achieved through multiple mechanisms, at the DNA, mRNA and protein levels. Our results also suggest RIT1 may be a promising therapeutic target for glioblastoma in future.

## MATERIALS AND METHODS

### Antibodies and chemicals

Antibodies and chemicals are listed in **Supplementary Table 1**.

### Cell culture, transfection and lentivirus infection

293T, HeLa cells and human malignant glioblastoma cell lines (U87-MG, U118-MG) were obtained from the American Type Culture Collection (ATCC). All cells were maintained in DMEM with 10% (v/v) FBS, and were grown at 37 °C with 5% CO_2_. Cells were transiently transfected using Lipofectamine 3000 (for plasmids transfection) or Lipofectamine RNAi MAX (for siRNA transfection) (Thermo, USA) according to the manufacturer’s instructions.

### Plasmid constructions

The plasmids used in this study were listed **Supplementary Table 1**. LZTR1 and RIT1 mutants were generated with the KOD-Plus Mutagenesis Kit (Toyobo, Japan) following the manufacturer’s instructions.

### RNA interference

Control siRNA and gene-specific ONTARGET SMARTpool siRNA pools for RBX1, CUL1-5 were purchased from Dharmacon (Thermo, USA), and the siRNAs for LZTR1 were purchased from GenePharma (China). Oligo sequences are listed in **Supplementary Table 2**.

### CRISP-Cas9 mediated gene knock out stable cell generation

pX459 plasmid (Addgene, USA) was used to clone guide oligos targeting LZTR1 or RIT1 gene. U87-MG cells were plated and transfected with pX459 constructs using Lipofectamine 3000 overnight. 24 hr after transfection, 1 μg/mL puromycin was used to screen cells for 3 days. Living cells were seeded in 96 well plate by limited dilution to isolate monoclonal cell line. Valid knock out cell lines are screened by corresponding antibody. Sequences of gene-specific sgRNAs are listed in **Supplementary Table 2**.

### Mass spectrometry analysis

HA-FLAG-LZTR1 complex was prepared by transfecting HA-FLAG-LZTR1 in 293T cells (5-10X100 mm dish). After 48 hr, the cells were lysed in RIPA buffer and the transfected LZTR1 was immunopurified from cell lysates with anti-FLAG M2 affinity gel (Sigma, USA) before being resolved by 10% SDS-PAGE. After Coomassie blue staining, the band corresponding to HA-FLAG-LZTR1 was excised. The liquid chromatograph tandem mass spectrometry analysis was carried out at the Proteomics Center of our institute.

### Cell proliferation assay

Cell proliferation rate was determined using Cell Counting Kit-8 (CCK-8) according to the manufacturer’s protocol (Dojindo Laboratories, Japan). Briefly, the cells were seeded onto 96-well plates at a density of 1200 cells per well. During a 4 to 7-d culture periods, 10 μL of the CCK-8 solution was added to cell culture, and incubated for 1 h. The resulting color was assayed at 450 nm using a microplate absorbance reader (Bio-Rad, USA). Each assay was carried out in triplicate.

### Cell cycle analysis

For cell cycle analysis, cells were harvested and washed 48 hr post-treatment with PBS, followed by propidium iodide (50 μg/mL) staining in the presence of RNase (10 μg/mL) for 30 minutes at 4 °C in the dark. The fraction of viable cells in G0/G1, S and G2/M phases of cell cycle were determined using a FACs flow cytometer and Cell Quest FACS system (Becton-Dickinson, USA). Results are representatives of three independent experiments with triples samples for each condition.

### EdU assay

EdU assay was performed using Cell-Light™ EdU Apollo®567 In Vitro Imaging Kit (Rib-bio, China). Treated and control cells (5 × 10^3^/well) were seeded onto 96-well plates and incubated with 5-ethynyl-20-deoxyuridine (EdU; 50 μM) for 2 hr at 37°C. Cells were fixed with 4% paraformaldehyde for 20 min, treated with 0.5% TritonX-100 for 10 min, washed with PBS for three times, and incubated with 100 μL of 1 X Apollo reagent for 30 min. Nuclei were labeled with Hochest 33342. The percentage of EdU-positive cells was calculated by image J software. Data reported represent the average of three independent experiments.

### Migration assay

Cell migration was determined by Transwell (Corning, USA) migration assay. U87-MG cell were precultured in serum-free medium for 48 hr. 1−3×10^4^ cells were seeded in serum-free medium in the upper chamber, and the lower chamber was filled with DMEM containing 10% FBS. After 48 h, the non-migrating cells on the upper chambers were carefully removed with a cotton swab, and the migrated cells underside of the filter stained and counted in nine different fields.

### Immunofluorescence and confocal microscopy

For immunofluorescence, cells were plated on chamber slides, fixed with 4% paraformaldehyde at room temperature for 30 min. After washing with PBS, cells were permeabilized with 0.2% Triton X-100 in PBS at room temperature for 5-10 min. Cells were then washed with PBS, blocked with 2% BSA in PBS for 1 hr, and incubated with primary antibodies in PBS for at 4 °C for overnight. After washing with PBS, fluorescence-labelled secondary antibodies were applied and DAPI was counterstained for 1 hr at room temperature. Cells were visualized and imaged using a confocal microscope (LSM710, Zeiss, Germany).

### *In vivo* ubiquitination assay

293T cells were transfected with HA–ubiquitin and the indicated constructs. 36 h after transfection, cells were treated with 30 μM MG132 for 6 h and then lysed in 1% SDS buffer (Tris [pH 7.5], 0.5 mM EDTA, 1 mM DTT) and boiled for 10 min. For immunoprecipitation, the cell lysates were diluted 10-fold in Tris-HCl buffer and incubated with anti-FLAG M2 agarose beads (Sigma) for 4 h at 4 °C. The bound beads are then washed four times with BC100 buffer (20 mM Tris-Cl, pH 7.9, 100 mM NaCl, 0.2 mM EDTA, 20% glycerol) containing 0.2% Triton X-100. The protein was eluted with 3X FLAG peptide for 2 h at 4 °C. The ubiquitinated form of RIT1 was detected by Western blot using anti-HA antibody.

### qRT-PCR assay

Total RNA was isolated from cells or tissues using TRIzol reagent (Thermo, USA), and cDNA was reverse-transcribed using the Superscript RT kit (Toyobo, Japan), according to the manufacturer’s instructions. Quantitative real-time PCR amplification was performed using the THUNDERBIRD SYBR qPCR Mix on ABI PRISM 7900HT instruments (Thermo, USA). The amplification was done in a total volume of 10 μL with the following steps: an initial denaturation step at 95 °C for 5 min, followed by 40 cycles for denaturation at 95°C for 15 sec and elongation at 60 °C for 45 sec. A melting curve analysis of each sample was used to check the specificity of amplification, and each sample was assayed in triplicate. GAPDH was used as the endogenous control, and the 2^−ΔΔCt^ method was used as relative quantification measure of differential expression. The primer sequences for qRT-PCR are listed in **Supplementary Table 2**.

### Generation of glioblastoma xenografts in mice

For intracranial xenograft studies, 6-to 9-week-old female nude mice (SLAC laboratory, China) were maintained in a barrier facility on high-efficiency particulate air (HEPA)-filtered racks. Luciferase-labeled U87-MG cells (1×10^6^) infected with LZTR1, RIT1 overexpression or deletion lentiviruses and their corresponding control cells were surgically implanted into left amygdala of mice brains using a stereotactic apparatus. For in vivo BLI, luciferase signals were detected 14, 20, 26 days after transplantation. Mice were intraperitoneally injected with D-luciferin (150mg/kg) (Promega, USA) and anesthetized with pentobarbital sodium. After 10 min of substrate administration, images were acquired through the In Vivo Imaging System. After transplantation, animals were closely followed and euthanized by cervical dislocation when they exhibited central nervous system symptoms, or drastic loss of body weight. Tumors were excised, formalin-fixed, paraffin-embedded, and sectioned for hematoxylin and eosin (HE) staining and IHC. All animal experiments were conducted with the approval of the Institutional Animal Care and Use Committee of Fudan University. All animal study was performed according to the Ethics Committee guidelines of Fudan University.

### Statistical analysis

All data are shown as mean values ± SD for experiments performed with at least three replicates. The difference between 2 groups was analyzed using paired Student’s t-test unless otherwise specified. Analysis of survival was conducted by Kaplan-Meier survival. * represents p < 0.05; ** represents p < 0.01; *** represents p < 0.001, ns represents not significant.

## ACKNOWLEDGMENTS

We sincerely thank Dr. Xiao Song (Department of Thoracic Surgery, Shanghai Pulmonary Hospital, Tongji University), Dr. Hexige Saiyin, Dr. Haoming Chen and Mrs.Taishan Min (School of Life Sciences, Fudan University) for their support. This work was supported by the National Natural Science Foundation of China (81972396, 81672558 and 81201533 to C.W.; 81572768, 81872260 to P.Z;81772657 to H.C.)

## AUTHOR CONTRIBUTIONS

D.L., C.W. designed and supervised the project. Y.W., J.Z., P.Z., Z.Z., Q.H., and D.Y. performed the experiments and data analyses. Y.W., J.Z., P.Z., Z.Z., Q.H. analyzed and interpreted the data. Y.W. wrote the manuscript. C.W. revised the manuscript. All authors read and approved the final manuscript.

## ABBREVIATIONS

LZTR1: leucine-zipper-like transcription regulator 1
RIT1: RAS-like without CAAX1
ATCC: American Type Culture Collection
siRNA: small interfering RNA
shRNA: short hairpin RNA
CCK-8: Cell Counting Kit-8
qRT-PCR: quantitative real-time polymerase chain reaction
BLI: bioluminescence imaging
IP: immunoprecipitation
DN: dominant negative
OS: overall survival
PFS: progression-free survival
TCGA: the cancer genome atlas
CGGA: Chinese glioma genome atlas
GBM: glioblastoma

## COMPLIANCE WITH ETHICS GUIDELINES

All institutional and national guidelines for the care and use of laboratory animals were followed. Yuqi Wang, Jianong Zhang, Pingzhao Zhang, Zhipeng Zhao, Qilin Huang, Dapeng Yun, Juxiang Chen, Hongyan Chen, Chenji Wang and Daru Lu declare that they have no conflict of interest.

## REFERENCES

1. Dolecek TA, Propp JM, Stroup NE, Kruchko C. CBTRUS statistical report: primary brain and central nervous system tumors diagnosed in the United States in 2005-2009. Neuro Oncol. 2012;14 Suppl 5:v1–49.

2. Dunn GP, Rinne ML, Wykosky J, Genovese G, Quayle SN, Dunn IF, et al. Emerging insights into the molecular and cellular basis of glioblastoma. Genes Dev. 2012;26(8):756–84.

3. Lee E, Yong RL, Paddison P, Zhu J. Comparison of glioblastoma (GBM) molecular classification methods. Semin Cancer Biol. 2018;53:201–11.

4. Liang Y, Diehn M, Watson N, Bollen AW, Aldape KD, Nicholas MK, et al. Gene expression profiling reveals molecularly and clinically distinct subtypes of glioblastoma multiforme. P Natl Acad Sci USA. 2005;102(16):5814–9.

5. Mischel PS, Nelson SF, Cloughesy TF. Molecular analysis of glioblastoma: pathway profiling and its implications for patient therapy. Cancer Biol Ther. 2003;2(3):242–7.

6. Diehn M, Nardini C, Wang DS, McGovern S, Jayaraman M, Liang Y, et al. Identification of noninvasive imaging surrogates for brain tumor gene-expression modules. Proc Natl Acad Sci U S A. 2008;105(13):5213–8.

7. Phillips HS, Kharbanda S, Chen R, Forrest WF, Soriano RH, Wu TD, et al. Molecular subclasses of high-grade glioma predict prognosis, delineate a pattern of disease progression, and resemble stages in neurogenesis. Cancer Cell. 2006;9(3):157–73.

8. Stupp R, Mason WP, van den Bent MJ, Weller M, Fisher B, Taphoorn MJ, et al. Radiotherapy plus concomitant and adjuvant temozolomide for glioblastoma. N Engl J Med. 2005;352(10):987–96.

9. Mischel PS, Cloughesy TF. Targeted molecular therapy of GBM. Brain Pathol. 2003;13(1):52–61.

10. Porter KR, McCarthy BJ, Freels S, Kim Y, Davis FG. Prevalence estimates for primary brain tumors in the United States by age, gender, behavior, and histology. Neuro Oncol. 2010;12(6):520–7.

11. Balca-Silva J, Matias D, Carmo AD, Sarmento-Ribeiro AB, Lopes MC, Moura-Neto V. Cellular and molecular mechanisms of glioblastoma malignancy: Implications in resistance and therapeutic strategies. Semin Cancer Biol. 2018.

12. Cancer Genome Atlas Research N. Comprehensive genomic characterization defines human glioblastoma genes and core pathways. Nature. 2008;455(7216):1061–8.

13. Broniscer A, Baker SJ, West AN, Fraser MM, Proko E, Kocak M, et al. Clinical and molecular characteristics of malignant transformation of low-grade glioma in children. J Clin Oncol. 2007;25(6):682–9.

14. Verhaak RG, Hoadley KA, Purdom E, Wang V, Qi Y, Wilkerson MD, et al. Integrated genomic analysis identifies clinically relevant subtypes of glioblastoma characterized by abnormalities in PDGFRA, IDH1, EGFR, and NF1. Cancer Cell. 2010;17(1):98–110.

15. Frattini V, Trifonov V, Chan JM, Castano A, Lia M, Abate F, et al. The integrated landscape of driver genomic alterations in glioblastoma. Nat Genet. 2013;45(10):1141–9.

16. Nacak TG, Leptien K, Fellner D, Augustin HG, Kroll J. The BTB-kelch protein LZTR-1 is a novel Golgi protein that is degraded upon induction of apoptosis. J Biol Chem. 2006;281(8):5065–71.

17. Stogios PJ, Prive GG. The BACK domain in BTB-kelch proteins. Trends Biochem Sci. 2004;29(12):634–7.

18. Johnston JJ, van der Smagt JJ, Rosenfeld JA, Pagnamenta AT, Alswaid A, Baker EH, et al. Autosomal recessive Noonan syndrome associated with biallelic LZTR1 variants. Genet Med. 2018;20(10):1175–85.

19. Umeki I, Niihori T, Abe T, Kanno SI, Okamoto N, Mizuno S, et al. Delineation of LZTR1 mutation-positive patients with Noonan syndrome and identification of LZTR1 binding to RAF1-PPP1CB complexes. Hum Genet. 2019;138(1):21–35.

20. Motta M, Fidan M, Bellacchio E, Pantaleoni F, Schneider-Heieck K, Coppola S, et al. Dominant Noonan syndrome-causing LZTR1 mutations specifically affect the Kelch domain substrate-recognition surface and enhance RAS-MAPK signaling. Hum Mol Genet. 2019;28(6):1007–22.

21. Deiller C, Van-Gils J, Zordan C, Tinat J, Loiseau H, Fabre T, et al. Coexistence of schwannomatosis and glioblastoma in two families. Eur J Med Genet. 2019:103680.

22. Piotrowski A, Xie J, Liu YF, Poplawski AB, Gomes AR, Madanecki P, et al. Germline loss-of-function mutations in LZTR1 predispose to an inherited disorder of multiple schwannomas. Nat Genet. 2014;46(2):182–7.

23. Evans DG, Bowers NL, Tobi S, Hartley C, Wallace AJ, King AT, et al. Schwannomatosis: a genetic and epidemiological study. J Neurol Neurosurg Psychiatry. 2018;89(11):1215–9.

24. Smith MJ, Isidor B, Beetz C, Williams SG, Bhaskar SS, Richer W, et al. Mutations in LZTR1 add to the complex heterogeneity of schwannomatosis. Neurology. 2015;84(2):141–7.

25. Shi GX, Cai W, Andres DA. Rit subfamily small GTPases: regulators in neuronal differentiation and survival. Cell Signal. 2013;25(10):2060–8.

26. Lein PJ, Guo X, Shi GX, Moholt-Siebert M, Bruun D, Andres DA. The novel GTPase Rit differentially regulates axonal and dendritic growth. J Neurosci. 2007;27(17):4725–36.

27. Cai W, Rudolph JL, Harrison SM, Jin L, Frantz AL, Harrison DA, et al. An evolutionarily conserved Rit GTPase-p38 MAPK signaling pathway mediates oxidative stress resistance. Mol Biol Cell. 2011;22(17):3231–41.

28. Shi GX, Andres DA. Rit contributes to nerve growth factor-induced neuronal differentiation via activation of B-Raf-extracellular signal-regulated kinase and p38 mitogen-activated protein kinase cascades. Mol Cell Biol. 2005;25(2):830–46.

29. Shi GX, Han J, Andres DA. Rin GTPase couples nerve growth factor signaling to p38 and b-Raf/ERK pathways to promote neuronal differentiation. J Biol Chem. 2005;280(45):37599–609.

30. Rusyn EV, Reynolds ER, Shao H, Grana TM, Chan TO, Andres DA, et al. Rit, a non-lipid-modified Ras-related protein, transforms NIH3T3 cells without activating the ERK, JNK, p38 MAPK or PI3K/Akt pathways. Oncogene. 2000;19(41):4685–94.

31. Feng YF, Lei YY, Lu JB, Xi SY, Zhang Y, Huang QT, et al. RIT1 suppresses esophageal squamous cell carcinoma growth and metastasis and predicts good prognosis. Cell Death Dis. 2018;9(11):1085.

32. Xu F, Sun S, Yan S, Guo H, Dai M, Teng Y. Elevated expression of RIT1 correlates with poor prognosis in endometrial cancer. Int J Clin Exp Pathol. 2015;8(9):10315–24.

33. Lawrence MS, Stojanov P, Mermel CH, Robinson JT, Garraway LA, Golub TR, et al. Discovery and saturation analysis of cancer genes across 21 tumour types. Nature. 2014;505(7484):495–501.

34. Forbes SA, Beare D, Gunasekaran P, Leung K, Bindal N, Boutselakis H, et al. COSMIC: exploring the world’s knowledge of somatic mutations in human cancer. Nucleic Acids Res. 2015;43(Database issue):D805–11.

35. Gomez-Segui I, Makishima H, Jerez A, Yoshida K, Przychodzen B, Miyano S, et al. Novel recurrent mutations in the RAS-like GTP-binding gene RIT1 in myeloid malignancies. Leukemia. 2013;27(9):1943–6.

36. Li JT, Liu W, Kuang ZH, Zhang RH, Chen HK, Feng QS. [Mutation and amplification of RIT1 gene in hepatocellular carcinoma]. Zhonghua Yi Xue Yi Chuan Xue Za Zhi. 2004;21(1):43–6.

37. Aoki Y, Niihori T, Banjo T, Okamoto N, Mizuno S, Kurosawa K, et al. Gain-of-function mutations in RIT1 cause Noonan syndrome, a RAS/MAPK pathway syndrome. Am J Hum Genet. 2013;93(1):173–80.

38. Gos M, Fahiminiya S, Poznanski J, Klapecki J, Obersztyn E, Piotrowicz M, et al. Contribution of RIT1 mutations to the pathogenesis of Noonan syndrome: four new cases and further evidence of heterogeneity. Am J Med Genet A. 2014;164A(9):2310–6.

39. Koenighofer M, Hung CY, McCauley JL, Dallman J, Back EJ, Mihalek I, et al. Mutations in RIT1 cause Noonan syndrome - additional functional evidence and expanding the clinical phenotype. Clin Genet. 2016;89(3):359–66.

40. Bertola DR, Yamamoto GL, Almeida TF, Buscarilli M, Jorge AA, Malaquias AC, et al. Further evidence of the importance of RIT1 in Noonan syndrome. Am J Med Genet A. 2014;164A(11):2952–7.

41. Chen PC, Yin J, Yu HW, Yuan T, Fernandez M, Yung CK, et al. Next-generation sequencing identifies rare variants associated with Noonan syndrome. Proc Natl Acad Sci U S A. 2014;111(31):11473–8.

42. Yamamoto GL, Aguena M, Gos M, Hung C, Pilch J, Fahiminiya S, et al. Rare variants in SOS2 and LZTR1 are associated with Noonan syndrome. J Med Genet. 2015;52(6):413–21.

43. Grobner SN, Worst BC, Weischenfeldt J, Buchhalter I, Kleinheinz K, Rudneva VA, et al. The landscape of genomic alterations across childhood cancers. Nature. 2018;555(7696):321–7.

44. Cancer Genome Atlas Research Network. Electronic address wbe, Cancer Genome Atlas Research N. Comprehensive and Integrative Genomic Characterization of Hepatocellular Carcinoma. Cell. 2017;169(7):1327–41 e23.

45. Bigenzahn JW, Collu GM, Kartnig F, Pieraks M, Vladimer GI, Heinz LX, et al. LZTR1 is a regulator of RAS ubiquitination and signaling. Science. 2018;362(6419):1171–7.

46. Steklov M, Pandolfi S, Baietti MF, Batiuk A, Carai P, Najm P, et al. Mutations in LZTR1 drive human disease by dysregulating RAS ubiquitination. Science. 2018;362(6419):1177–82.

47. Castel P, Cheng A, Cuevas-Navarro A, Everman DB, Papageorge AG, Simanshu DK, et al. RIT1 oncoproteins escape LZTR1-mediated proteolysis. Science. 2019;363(6432):1226–30.

