## Supplementary material for "LZTR1 inactivation promotes MAPK/ ERK pathway activation in glioblastoma by stabilizing oncoprotein RIT1"

**Supplementary materials**

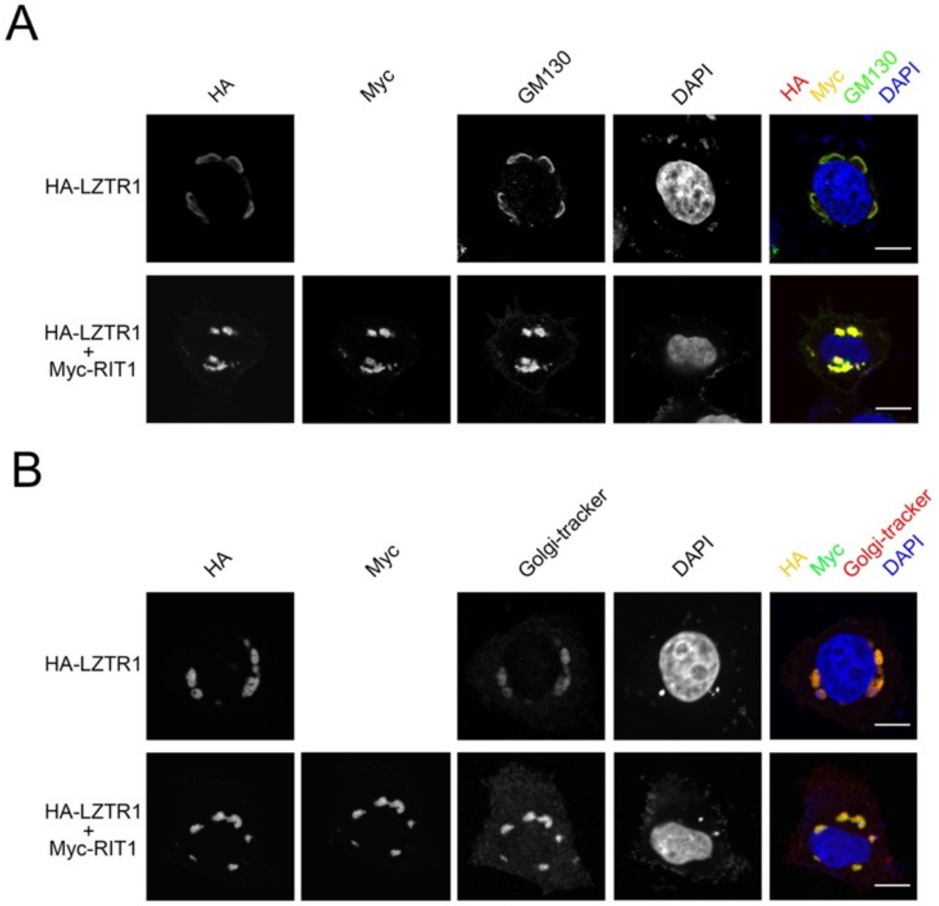


**Supplementary Figure 1. LZTR1 recruits RIT1 to Golgi apparatus.** (A) Representative immunofluorescence images of HeLa cells transfected with indicated plasmids, stained with LZTR1(HA), RIT1(Myc), GM130 and DAPI. Scale bar, 20 μm. (B) Representative immunofluorescence images of HeLa cells transfected with indicated plasmids, stained with LZTR1(HA), RIT1(Myc), Golgi-tracker Red dyes and DAPI. Scale bar, 20 μm.

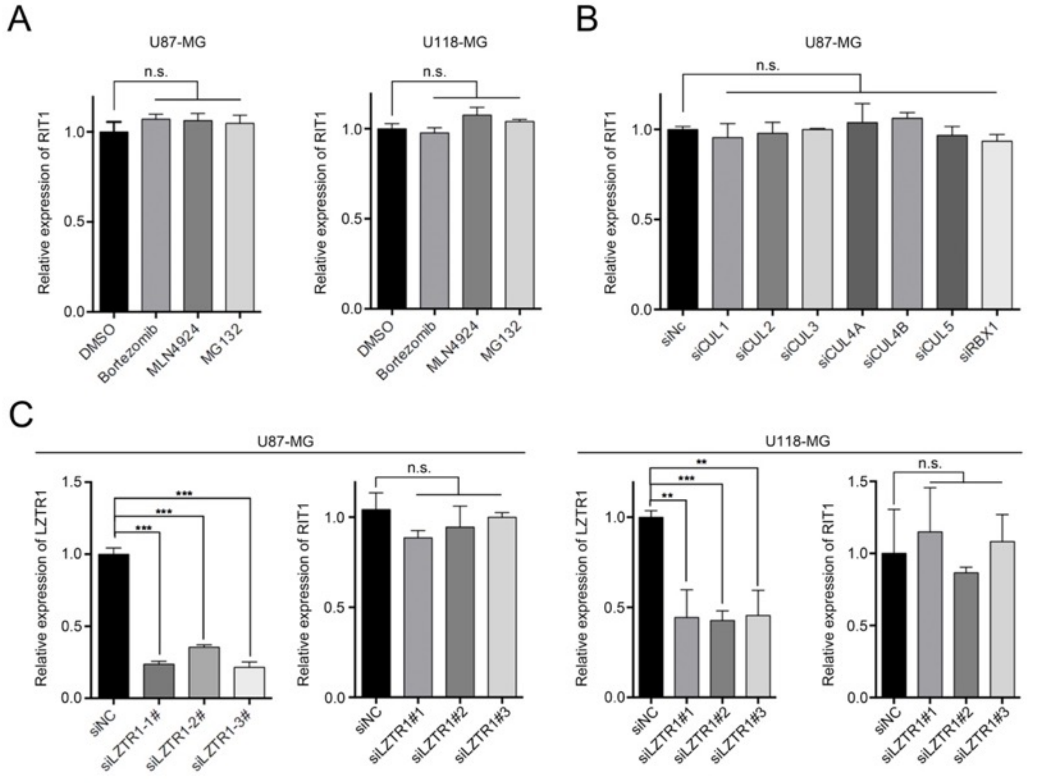


**Supplementary Figure 2. LZTR1 does not regulate RIT1 mRNA level.** (A) qRT-PCR assessment of RIT1 mRNA expression in U87-MG and U118-MG cells treated with DMSO, MG132 (20 μM), Bortezomib (20 nM) or with MLN4924 (100 nM) for 8 hr. Data are shown as means ± SD(n=3). (B) qRT-PCR assessment of RIT1 mRNA expression in U87-MG cells transfected with indicated siRNAs. Data are shown as means ± SD(n=3). (C) qRT-PCR assessment of LZTR1 or RIT1 mRNA expression in U87-MG (left panel) and U118-MG (right panel) cells transfected with indicated siRNAs. Data are shown as means ± SD(n=3). **$p<0.01$, ***$p<0.001$.

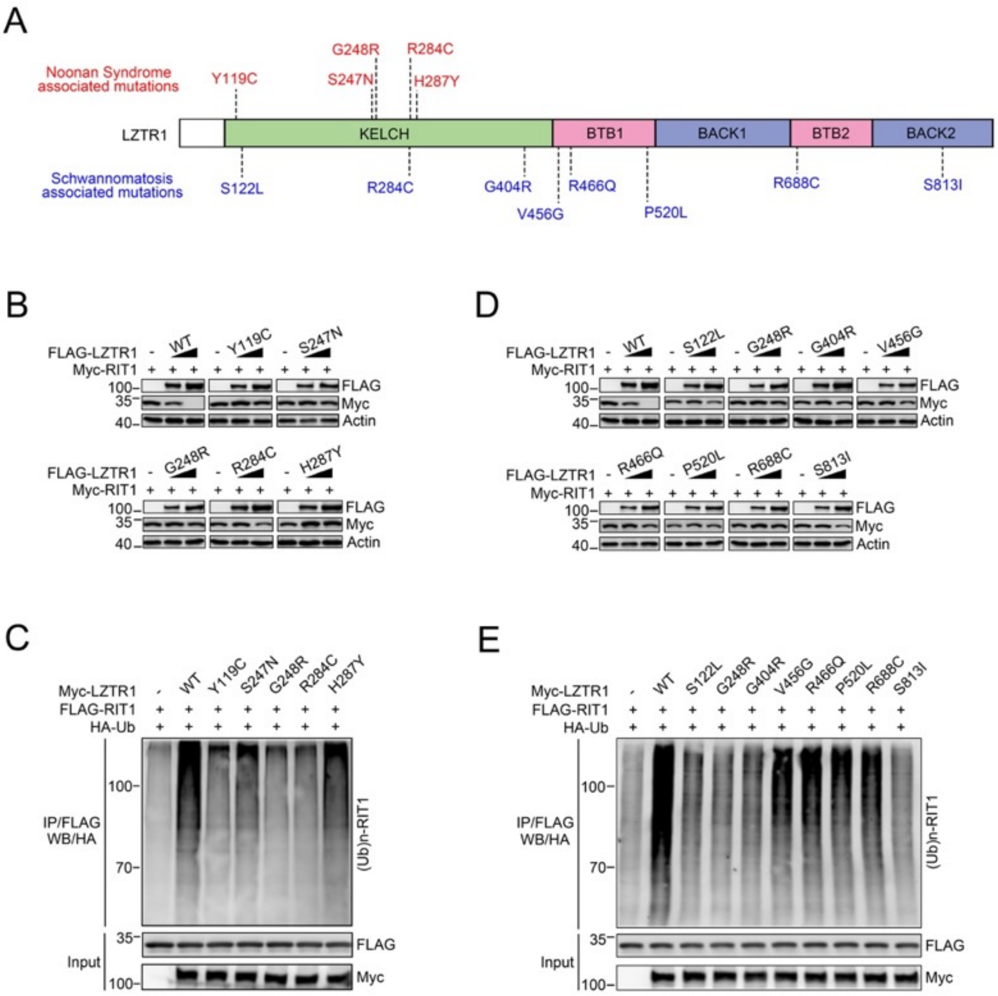


**Supplementary Figure 3. Noonan syndrome and schwannoma-associated of LZTR1 mutants are defective in promoting RIT1 degradation and ubiquitination.** (A) Diagram showing the Noonan syndrome and schwannoma-associated LZTR1 mutations. (B) Western blot of indicated proteins in WCLs from 293T cells transfected with the indicated plasmids. Noonan syndrome-associated of LZTR1 mutants were used. (C) Western blot of the products of in vivo ubiquitination assays from 293T cells infected with the indicated plasmids and treated with 20 μM MG132 for 8 h. Noonan syndrome-associated of LZTR1 mutants were used. (D) Western blot of indicated proteins in WCLs from 293T cells transfected with the indicated plasmids. schwannoma-associated of LZTR1 mutants were used. (E) Western blot of the products of in vivo ubiquitination assays from 293T cells infected with the indicated plasmids and treated with 20 μM MG132 for 8 h. schwannoma-associated of LZTR1 mutants were used.

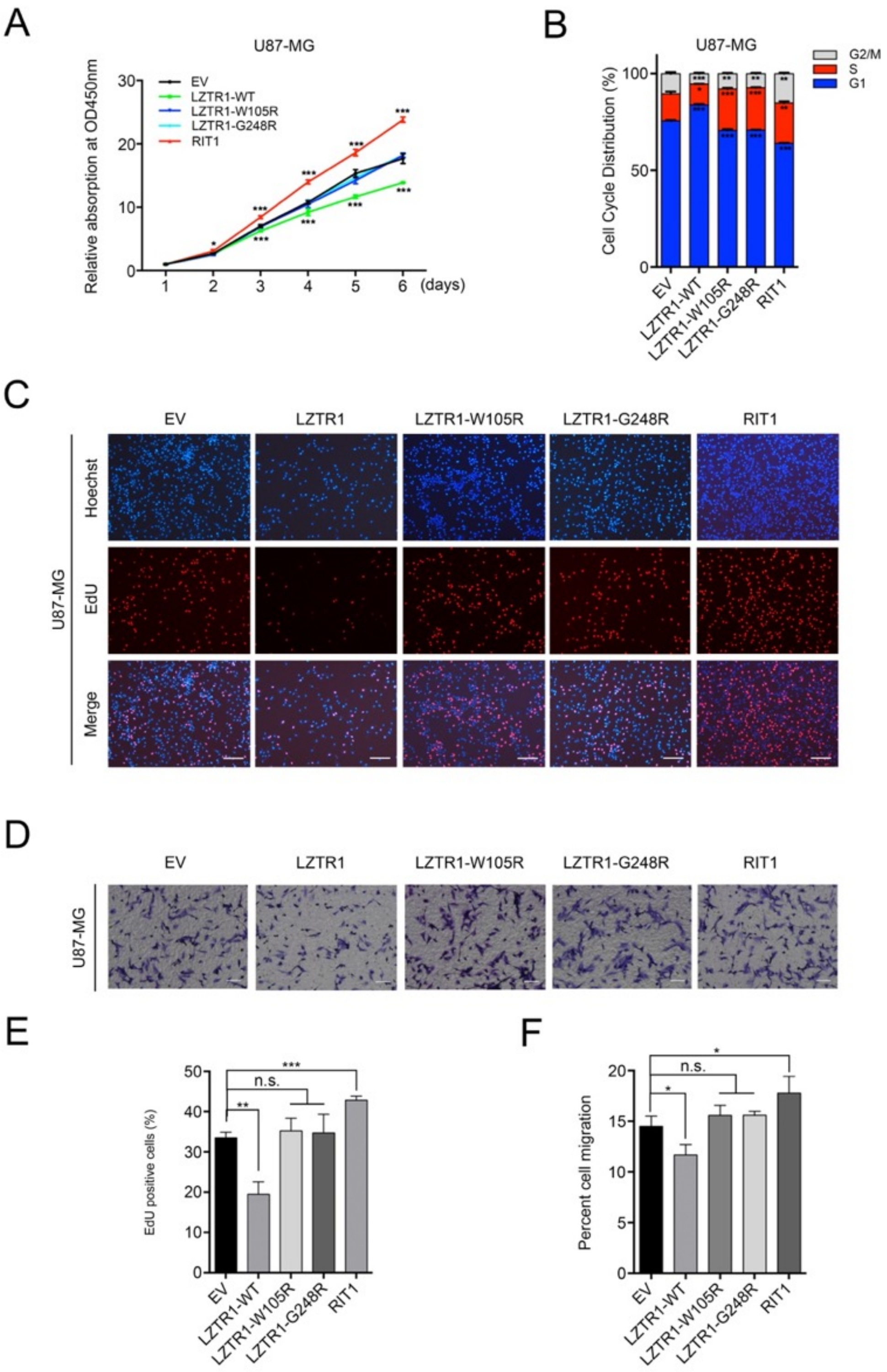


**Supplementary Figure 4. LZTR1 and RIT1 regulated cell growth and migration.** (A) CCK-8 cell proliferation analysis of U87-MG cells infected with lentivirus expressing indicated proteins. Data are shown as mean ± SD (n=3). ***p<0.001. (B) Cell cycle analysis of U87-MG cells infected with lentivirus expressing indicated proteins. Data are shown as means ± SD (n=3). *p<0.05, **p<0.01, ***p<0.001. (C) EdU incorporation analysis of U87-MG cells infected with lentivirus expression indicated proteins. (D) Cell migration analysis of U87-MG cells infected with lentivirus expression indicated proteins. Scale bar, 20 μm. (E, F) The quantitative analysis of EdU incorporation assay (E), and cell migration assay (F). Data are shown as means ± SD (n=3). *p<0.05, **p<0.01, ***p<0.001.

**Supplementary Table 1. Key resource table**

| REAGENT or RESOURCE | SOURCE | IDENTIFIER |
|---|---|---|
| Antibodies | | |
| α-LZTR1 | ABclonal | Cat#A7350 |
| α-RIT1 | ABGENT | Cat#AP14501a |
| α-His | AOGMA | Cat#9618 |
| α-FLAG | MBL | Cat#M185-7 |
| α-Myc | MBL | Cat#M192-7 |
| α-HA | MBL | Cat#M180-7 |
| α-$\beta$-cantenin | R&D | Cat#AF1329 |
| α-p21 | Proteintech | Cat#3733-1 |
| α-Actin | ABclonal | Cat#AC028 |
| α-CUL1 | Abcam | Cat#ab75817 |
| α-CUL2 | Proteintech | Cat#10981-2-AP |
| α-CUL3 | Proteintech | Cat#11107-1-AP |
| α-CUL4A | EPITOMICS | Cat#2715-S |
| α-CUL4B | Proteintech | Cat#12916-1-AP |
| α-CUL5 | Abcam | Cat#ab34840 |
| α-RBX1 | Proteintech | Cat#14895-1-AP |
| α-SOS1 | Abcam | Cat#ab140621 |
| α-SHP2 | Abcam | Cat#ab32083 |
| α-A-RAF | Abcam | Cat#ab200653 |
| α-B-RAF | Abcam | Cat#ab33899 |
| α-C-RAF | Abcam | Cat#ab181115 |
| α-MEK1 | Abcam | Cat#ab32091 |
| α-MEK2 | Abcam | Cat#ab32517 |
| α-ERK1/2 | Cell Signaling Technology | Cat#4695T |
| α-PAN-RAS | Abcam | Cat#ab52939 |
| α-R-RAS | Abcam | Cat#ab191399 |
| α-N-RAS | Proteintech | Cat#10724-1-AP |
| α-K-RAS | ABclonal | Cat#A1190 |
| α-H-RAS | Proteintech | Cat#18295-1-AP |
| α-MSK1 | Abcam | Cat#ab155405 |
| α-MEK1/2 | Cell Signaling Technology | Cat#4694 |
| α-pERK1/2 | Cell Signaling Technology | Cat#4370T |
| α-pMEK1/2 | Cell Signaling Technology | Cat#3958 |
| α-pMSK1 | Abcam | Cat#ab79499 |
| Bacterial and Virus Strains | | |
| *E. coli* DH5α | This paper | N/A |

| | | |
|---|---|---|
| Topo 10 competent *E. coli* | This paper | N/A |
| Rosetta (DE3) competent cells | This paper | N/A |
| Chemicals, Peptides, and Recombinant Proteins | | |
| Anti-FLAG M2 affinity gel | Sigma | Cat#A2220 |
| Protein A/G beads | Thermo Fisher | Cat#20421 |
| MG132 | Selleckchem | Cat#S2619 |
| Bortezomib | Selleckchem | Cat#S1013 |
| MLN4924 | Selleckchem | Cat#S7109 |
| CHX | MCE | Cat#HY-12320 |
| Puromycin | Sigma | Cat#P8833 |
| Trizol | Invitrogen | Cat#14496026 |
| Lipofectamine 2000 | Thermo Fisher | Cat#11668027 |
| Lipofectamine 3000 | Thermo Fisher | Cat#L3000008 |
| Lipofectamine RNAiMax | Thermo Fisher | Cat#13778030 |
| D-luciferin | Promega | Cat#E1603 |
| FLAG peptide | ChinaPeptides | Cat#04010006736 |
| GST-LZTR1 | This paper | N/A |
| GST-RIT1 | This paper | N/A |
| Critical Commercial Assays | | |
| Cell Counting Kit-8 | Dojindo Laboratories | Cat#CK04 |
| EdU Apollo 567 Cell Tracking Kit | Rib-bio | Cat#C10310-1 |
| Golgi-Tracker Red | Beyotime | Cat#C1403 |
| KOD-Plus-Mutagenesis Kit | Toyobo | Cat#SMK-101 |
| THUNDERBIRD SYBR qPCR Mix | Toyobo | Cat#QPS-201 |
| Experimental Models: Cell Lines | | |
| 293T | ATCC | Cat#CRL-3216 |
| HeLa | ATCC | Cat#CCL-2 |
| U87-MG | ATCC | Cat#HTB-14 |
| U118-MG | ATCC | Cat#HTB-15 |
| Experimental Models: Orgnisms/Strains | | |
| Mouse: Nude mice | SLAC laboratory animal Center | N/A |
| Recombinant DNA | | |
| pCMV-HA-FLAG-LZTR1 | This paper | N/A |
| pCMV-FLAG/Myc/HA-LZTR1 | This paper | N/A |
| pCMV-FLAG/Myc-RIT1 | This paper | N/A |
| pCMV-FLAG/Myc-KLHL8 | This paper | N/A |
| pCMV-FLAG/Myc-KLHL9 | This paper | N/A |
| pCMV-FLAG/Myc-KLHL11 | This paper | N/A |
| pCMV-FLAG/Myc-KLHL26 | This paper | N/A |
| pCMV-FLAG-CUL1-DN | Addgene | Cat#15818 |
| pCMV-FLAG-CUL2-DN | Addgene | Cat#15819 |

| | | |
|---|---|---|
| pCMV-FLAG-CUL3-DN | Addgene | Cat#15820 |
| pCMV-FLAG-CUL4A-DN | Addgene | Cat#15821 |
| pCMV-FLAG-CUL4B-DN | Addgene | Cat#15822 |
| pCMV-FLAG-CUL5-DN | Addgene | Cat#15823 |
| pCMV-FLAG-R-RAS | This paper | N/A |
| pCMV-FLAG-H-RAS | This paper | N/A |
| pCMV-FLAG-K-RAS | This paper | N/A |
| pCMV-FLAG-PTPN11 | This paper | N/A |
| pCMV-FLAG/Myc-LZTR1 (M1~M10) | This paper | N/A |
| pCMV-FLAG-RIT1 ($\Delta 1 \sim \Delta 10$) | This paper | N/A |
| pCMV-FLAG/Myc-LZTR1-W105R | This paper | N/A |
| pCMV-FLAG/Myc-LZTR1-D139A | This paper | N/A |
| pCMV-FLAG/Myc-LZTR1-N143T | This paper | N/A |
| pCMV-FLAG/Myc-LZTR1-G195S | This paper | N/A |
| pCMV-FLAG/Myc-LZTR1-R198G | This paper | N/A |
| pCMV-FLAG/Myc-LZTR1-G248R | This paper | N/A |
| pCMV-FLAG/Myc-LZTR1-R284S | This paper | N/A |
| pCMV-FLAG/Myc-LZTR1-T288I | This paper | N/A |
| pCMV-FLAG/Myc-LZTR1-G404E | This paper | N/A |
| pCMV-FLAG/Myc-LZTR1-R810W | This paper | N/A |
| pCMV-FLAG/Myc-LZTR1-Y119C | This paper | N/A |
| pCMV-FLAG/Myc-LZTR1-S247N | This paper | N/A |
| pCMV-FLAG/Myc-LZTR1-H287Y | This paper | N/A |
| pCMV-FLAG/Myc-LZTR1-S122L | This paper | N/A |
| pCMV-FLAG/Myc-LZTR1-G404R | This paper | N/A |
| pCMV-FLAG/Myc-LZTR1-V456G | This paper | N/A |
| pCMV-FLAG/Myc-LZTR1-R466Q | This paper | N/A |
| pCMV-FLAG/Myc-LZTR1-P520L | This paper | N/A |
| pCMV-FLAG/Myc-LZTR1-R688C | This paper | N/A |
| pCMV-FLAG/Myc-LZTR1-S813I | This paper | N/A |
| pCD513B-LZTR1 | This paper | N/A |
| pCD513B-RIT1 | This paper | N/A |
| pCD513B-LZTR1-W105R | This paper | N/A |
| pCD513B-LZTR1-G248R | This paper | N/A |
| GFP-TGNP | This paper | N/A |
| pGEX-4T-2-LZTR1 | This paper | N/A |
| His-RIT1 | This paper | N/A |
| pCMV-HA-Ub | This paper | N/A |
| Software | | |
| Prism 6 | Graphpad | http://www.graphpad.com/ |

**Supplementary Table 2. Primers used for RT-qPCR in cultured cell lines, and sequences**

**of shRNAs, siRNAs, and sgRNAs.**

| **Primers for RT-qPCR with cell lines samples** | | |
|---|---|---|
| Gene name | F: 5’-3’ | R: 5’-3’ |
| LZTR1 | GCGGTTCGATGTGAAAGACT | TGTAACCCCCAAAGACAAACA |
| RIT1 | TAGGGAAGAGTGCCATGACC | CAGGCTCATCATCAATACGG |
| GAPDH | GAAGGTGAAGGTCGGAGT | GAAGATGGTGATGGGATTTC |
| **Sequences of shRNAs** | | |
| Gene name | Sequence | |
| shLZTR1#1 | CGGGACAAGATGTTTGTAT | |
| shLZTR1#3 | GCGGAATTCTGTGACATCA | |
| shRIT1#2 | AGAATTCAGCTGTCCCTTT | |
| shRIT1#3 | TCGAAGTTTCCATGAAGTT | |
| Sequences of siRNAs | | |
| Gene name | Sequence | |
| siLZTR1#1 | GGACAUUUAUUCCAAUUCU | |
| siLZTR1#2 | CGGGACAAGAUGUUUGUAU | |
| siLZTR1#3 | GCGGAAUUCUGUGACAUCA | |
| siRBX1#1 | GAAGCGCUUUGAAGUGAAA | |
| siRBX1#2 | GGGAUAUUGUGGUUGAUAA | |
| siRBX1#3 | GGAACCACAUUAUGGAUCU | |
| siRBX1#4 | CAUAGAAUGUCAAGCUAAC | |
| siCUL1#1 | CAACGAAGAGUUCAGGUUU | |
| siCUL1#2 | CGAGGAAGACCGCAAACUA | |
| siCUL1#3 | AGACAGUGCUUGAUGUUCA | |
| siCUL1#4 | CAUAGAAGACAAAGACGUA | |
| siCUL2#1 | GGAAGUGCAUGGUAAAUUU | |
| siCUL2#2 | CAUCCAAGUUCAUAUACUA | |
| siCUL2#3 | GCAGAAAGACACACCACAA | |
| siCUL2#4 | UGGUUUACCUCAUAUGAUU | |
| siCUL3#1 | GAGAAGATGTACTAAATTC | |
| siCUL3#2 | CGACAGAAAACATGAGATA | |
| siCUL3#3 | GAAGTAGACGACGACAGA | |
| siCUL3#4 | GAGATCAAGTTGTACGTTA | |
| siCUL4A#1 | GCACAGAUCCUUCCGUUUA | |
| siCUL4A#2 | GAACAGCGAUCGUAAUCAA | |
| siCUL4A#3 | GCAUGUGGAUUCAAAGUUA | |
| siCUL4A#4 | GCGAGUACAUCAAGACUUU | |
| siCUL4B#1 | UAAAUAACCUCCUUGAUGA | |
| siCUL4B#2 | CAGAAGUCAUUAAUUGCUA | |
| siCUL4B#3 | CGGAAAGAGUGCAUCUGUA | |
| siCUL4B#4 | GCUAUUGGCCGACAUAUGU | |
| siCUL5#1 | GACACGACGUCUUAUAUUA | |
| siCUL5#2 | GCAAAUAGAGUGGCUAAUA | |

| | |
|---|---|
| siCUL5#3 | UAAACAAGCUUGCUAGAAU |
| siCUL5#4 | CGUCUAAUCUGUUAAAGAA |
| Sequences of sgRNAs | |
| Gene name | Sequence |
| sgLZTR1#1 | AGTCTTTCACATCGAACCGC |
| sgLZTR1#6 | GTCTCCACCAAATACATAAA |
| sgRIT1#2 | TCGGTGGCTGATGAACTGCA |
| sgRIT1#3 | TGATGATGAGCCTGCCAATC |
